# The *Streptococcus pneumoniae* transcriptome in patient cerebrospinal fluid identifies novel virulence factors required for meningitis

**DOI:** 10.1101/2024.04.30.591261

**Authors:** Emma C Wall, José Afonso Guerra-Assunção, Marie Yang, Rutger Koning, Teerawit Audshasai, Alizé Proust, Rieza Aprianto, Elisa Ramos-Sevilliano, Giuseppe Ercoli, Modupeh Betts, Nicola Bordin, Vanessa S. Terra, Jan-Willem Veening, David G Lalloo, Brendan W Wren, Robert J Wilkinson, Matthijs C Brouwer, Diederik van de Beek, Aras Kadioglu, Robert S Heyderman, Jeremy S Brown

## Abstract

To better understand *Streptococcus pneumoniae* pathogenesis we performed RNA sequencing on cerebrospinal fluid (CSF) from meningitis patients to identify bacterial genes expressed during invasion of the central nervous system. Comparison to transcriptome data for serotype 1 *S. pneumoniae* cultured in *ex vivo* human CSF defined a subset of 57 genes with high expression during human meningitis. Deletion of two of the most highly expressed genetic loci, *bgaA* (encodes for a ß-galactosidase) or the SP_1801-5 putative stress response operon, resulted in *S. pneumoniae* strains still able to transmigrate the blood brain barrier but which were more susceptible to complement opsonisation and unable to maintain brain infection in a murine meningitis model. In 1144 meningitis patients, infection with *bgaA* containing *S. pneumoniae* strains was associated with a higher mortality (22% versus 14% p=0.02). These data demonstrate that direct bacterial RNAseq from CSF can identify previously undescribed *S. pneumoniae* virulence factors required for meningitis pathogenesis.

## Introduction

Acute bacterial meningitis is a leading cause of infectious mortality and morbidity world-wide, with an estimated 2.8 million cases in 2016^1,2^. *Streptococcus pneumoniae* remains the most frequent cause in most regions, particularly in sub-Saharan Africa^3^ where mortality reaches 30-60% and survivors frequently experience long-term neurological sequelae^4,5^. In contrast to high income settings, the poor outcomes in sub-Saharan Africa are not improved by adjunctive therapies^6–9^, and an improved understanding of meningitis pathogenesis is needed to idenitfy novel therapeutic approaches^10^. Previous work has described how *S. pneumoniae* translocates across the blood brain barrier^11–13,14^ and causes neurotoxicity through the toxin pneumolysin (Ply), production of hydrogen peroxide, and the pro-inflammatory effects of cell wall components^15–19^. However, the bacterial factors required for *S. pneumoniae* invasion and growth in the central nervous system (CNS) during meningitis remain largely unknown. Animal model experiments, including a mutant library screen in rabbits^20^ and transcriptomic studies using mouse, zebrafish or rabbit models have identified multiple genes postulated to be important for meningitis pathogenesis, but how these data relate to human disease remains unclear^21–24^. To address this problem, we used RNAseq to profile the bacterial transcriptome in pre-antibiotic CSF samples from patients with pneumococcal meningitis to make a comprehensive assessment of *S. pneumoniae* gene expression during invasive human infection and provide a global overview of the bacterial adaptation to meningeal invasion. Cross-referencing *S. pneumoniae* gene expression under meningitis-like conditions *in vitro*, with bacterial transcriptomes from patients identifed bacterial genes uniquely highly expressed during human meningitis. From these comparisons we identified two highly expressed genetic loci (*bgaA* and Sp_1801-05) in patient CSF with no documented or hypothetical role in meningitis for further investigation.

## Results

### The S. pneumoniae meningitis transcriptome is dominated by genes involved in replication, metabolism and virulence

To improve understanding of *S. pneumoniae* adaptation during human meningitis, we isolated bacterial RNA from pre-antibiotic CSF samples obtained from Malawian adults subsequently confirmed to have *S. pneumoniae* meningitis by culture or PCR (Table 1). Of the 36 samples processed, quantifable RNA suitable for bacterial RNAseq was obtained and sequenced from 11 participants (median age 30.5 years, 46% male, 91% HIV seropositive, 10/11 (91%) died).^25^ Since the pneumococcal strains causing the meningitis were unknown, reads could not be mapped to a single unifying pneumococcal genome, and the data were first aligned to a collection of 70 curated and complete *S. pneumoniae* genomes. The alignment ratio between samples varied from 0.242 to 0.957 (mean 0.82), with one outlier sample (194C) with low RNA abundance showing relatively low alignment (**Table 1**, **Figure 1)**. Log_2_ values for maximum and minimum gene transcript/million reads (TPM) abundance ranged from 1.00 to 15.68 (**Supplemental data file 1**). Of the 70 *S. pneumoniae* strains screened, the highest serotype alignments for an individual sample were serotypes 1 (n=4 samples) and 3 (n=3 samples), similar to the known distribution of serotypes causing meningitis in Malawi^26,27^. We further analysed each sample individually against the corresponding best-matched *S. pneumoniae* strain and used normalized gene expression to identify between 30 to 118 highly expressed genes per sample (**Figure 1A**, listed in **Supplemental Table 4)**. Of these, only six genes were uniformly highly expressed in all 11 samples (*rnpB, ssrA, tuf*, *spxB*, *rpmE2*, *gap*). However, many other highly transcribed genes from individual samples overlapped across samples (**Figure 1B**). To optimize our analysis by re-mapping and aligning the RNAseq data to a unifying consensus strain, we selected the genome of the serotype 1 strain gamPNI0373 (genbank CP001845.1, isolated from a child with sepsis in Ghana) which showed the highest degree of alignment across all 11 transcriptome samples (76% to 93%) (**Table 1**). This generated a single table of genes aligned to an *S. pneumoniae* genome for subsequent analysis (**Table 1).** A heatmap of the top 250 most highly transcribed genes confirmed that the 50 highly expressed genes were conserved across multiple samples (**Figure 1C**). Using a threshold expression level of >10.62 log_2_ transcripts per million (TPM), 1.5+ SDs above the mean log_2_ TPM (6.89, SD 2.49), 102 genes were defined as highly transcribed in CSF across all samples and were ranked according to their median TPM (**Table 2)**.

**Figure 1:**
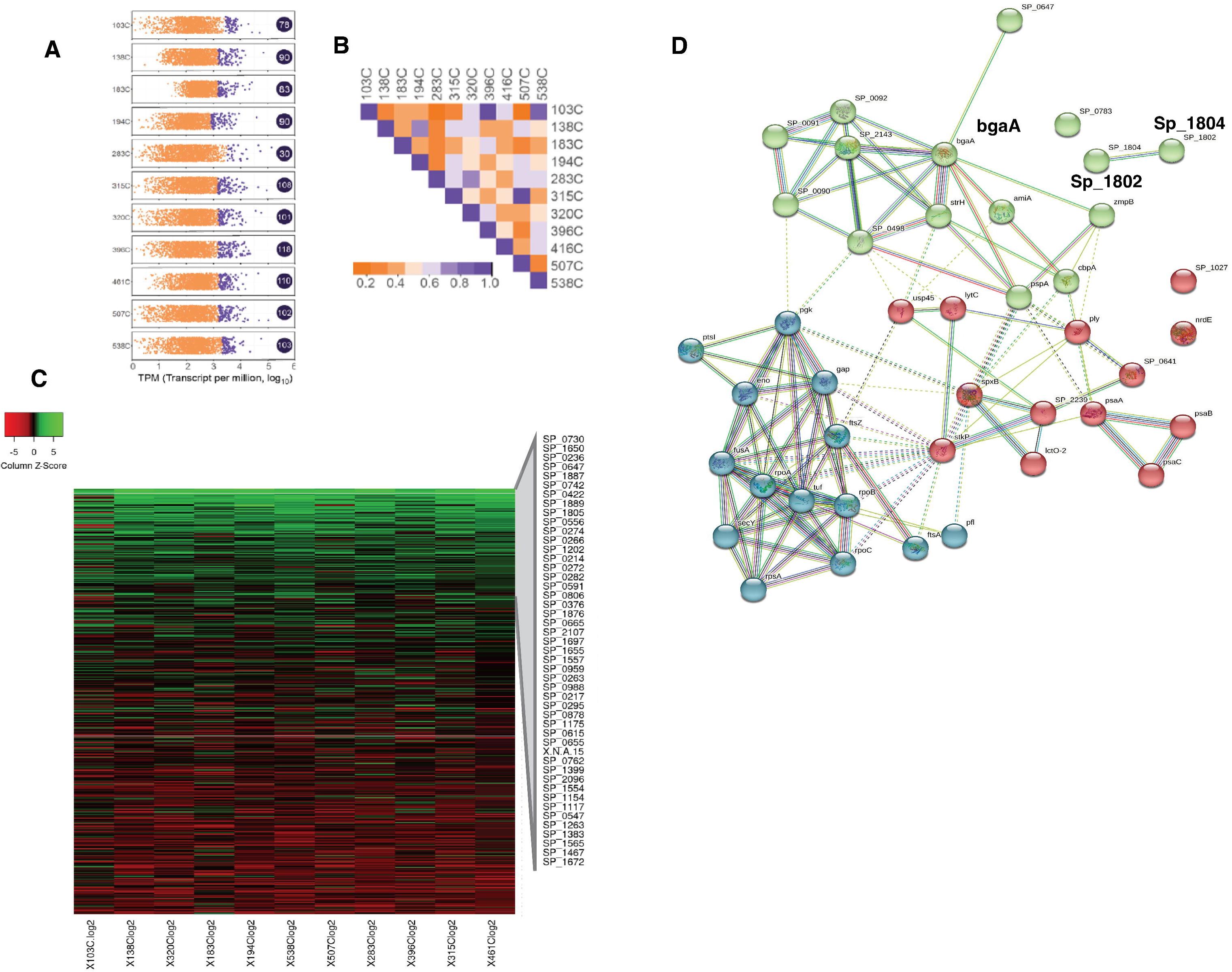
*Streptococcus pneumoniae* prioritises co-expression of metabolic, cell replication and virulence genes during infection of the CSF in meningitis. (A) Distribution of pneumococcal gene expression (log2 TPM) across 11 CSF samples from adults with meningitis. Each dot represents in individual gene, dots coloured purple represent highly expressed (>75th centile) genes. (B) Correlation matrix of gene expression across samples. (C) Heatmap showing the top 50 genes are consistently highly expressed across all human CSF samples. (D) 50 most highly expressed genes fall into three clusters (STRING Network Plot), annotated for cellular replication (blue), metabolism (green) and virulence (red). Each node represents an individual gene, the number of edges between nodes, and distance represents the strength of the association (co-expression).

**Table 1:**
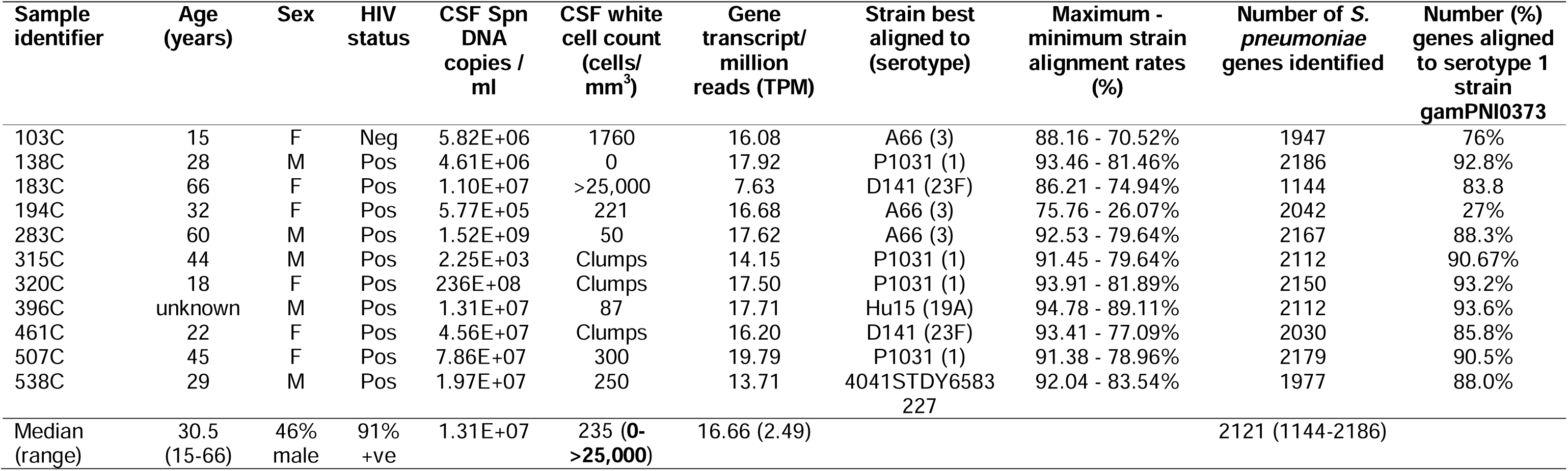
Summary of patient and sample characteristics.

| Sample identifier | Age (years) | Sex | HIV status | CSF Spn DNA copies / ml | CSF white cell count (cells/mm<sup>3</sup>) | Gene transcript/ million reads (TPM) | Strain best aligned to (serotype) | Maximum - minimum strain alignment rates (%) | Number of *S. pneumoniae* genes identified | Number (%) genes aligned to serotype 1 strain gamPNI0373 |
| --- | --- | --- | --- | --- | --- | --- | --- | --- | --- | --- |
| 103C | 15 | F | Neg | 5.82E+06 | 1760 | 16.08 | A66 (3) | 88.16 - 70.52% | 1947 | 76% |
| 138C | 28 | M | Pos | 4.61E+06 | 0 | 17.92 | P1031 (1) | 93.46 - 81.46% | 2186 | 92.8% |
| 183C | 66 | F | Pos | 1.10E+07 | >25,000 | 7.63 | D141 (23F) | 86.21 - 74.94% | 1144 | 83.8 |
| 194C | 32 | F | Pos | 5.77E+05 | 221 | 16.68 | A66 (3) | 75.76 - 26.07% | 2042 | 27% |
| 283C | 60 | M | Pos | 1.52E+09 | 50 | 17.62 | A66 (3) | 92.53 - 79.64% | 2167 | 88.3% |
| 315C | 44 | M | Pos | 2.25E+03 | Clumps | 14.15 | P1031 (1) | 91.45 - 79.64% | 2112 | 90.67% |
| 320C | 18 | F | Pos | 236E+08 | Clumps | 17.50 | P1031 (1) | 93.91 - 81.89% | 2150 | 93.2% |
| 396C | unknown | M | Pos | 1.31E+07 | 87 | 17.71 | Hu15 (19A) | 94.78 - 89.11% | 2112 | 93.6% |
| 461C | 22 | F | Pos | 4.56E+07 | Clumps | 16.20 | D141 (23F) | 93.41 - 77.09% | 2030 | 85.8% |
| 507C | 45 | F | Pos | 7.86E+07 | 300 | 19.79 | P1031 (1) | 91.38 - 78.96% | 2179 | 90.5% |
| 538C | 29 | M | Pos | 1.97E+07 | 250 | 13.71 | 4041STDY6583<br>227 | 92.04 - 83.54% | 1977 | 88.0% |
| Median (range) | 30.5 (15-66) | 46% male | 91% +ve | 1.31E+07 | 235 (0-<br>**>25,000**) | 16.66 (2.49) |  |  | 2121 (1144-2186) |  |

**Table 2:**
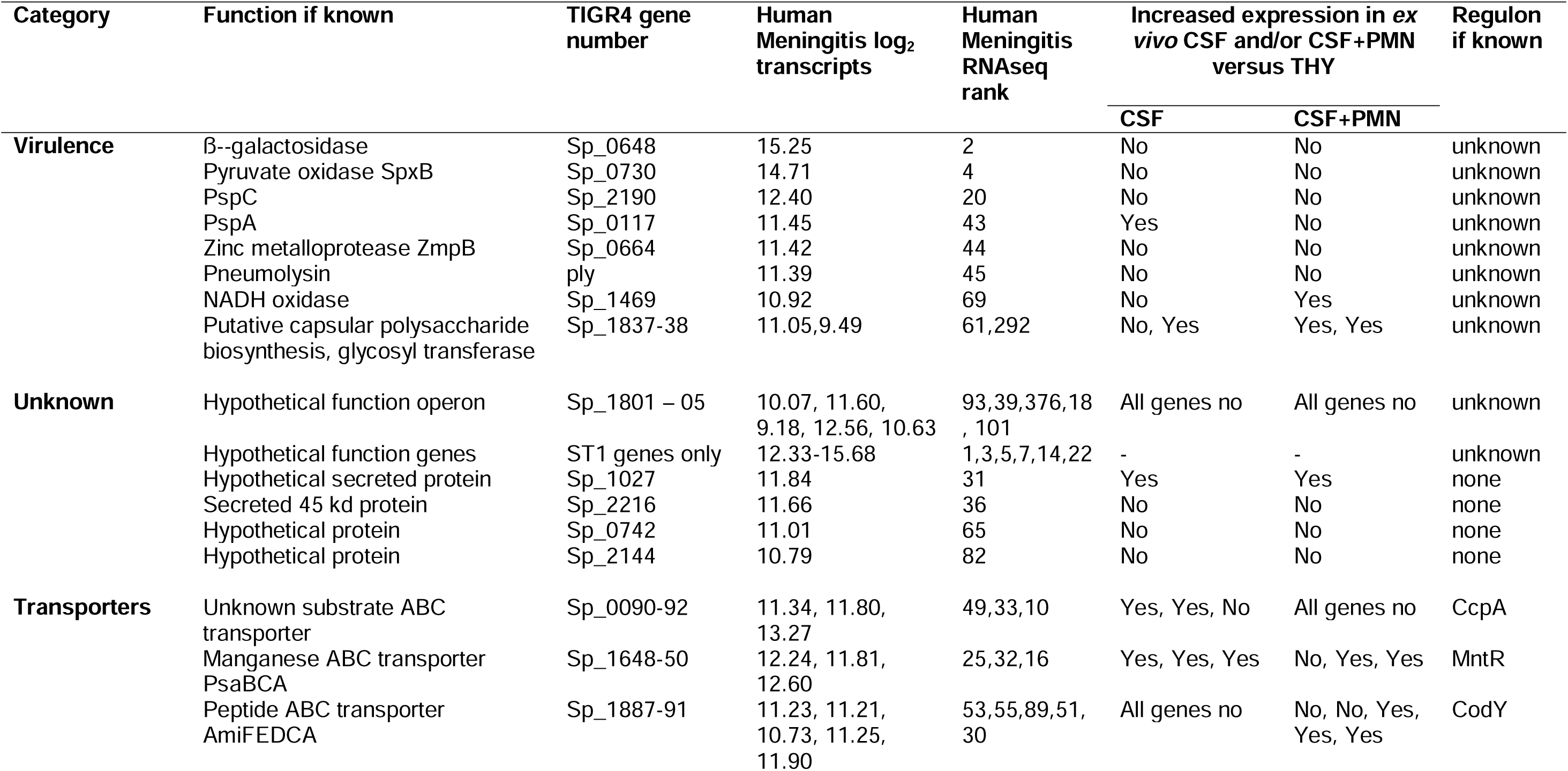
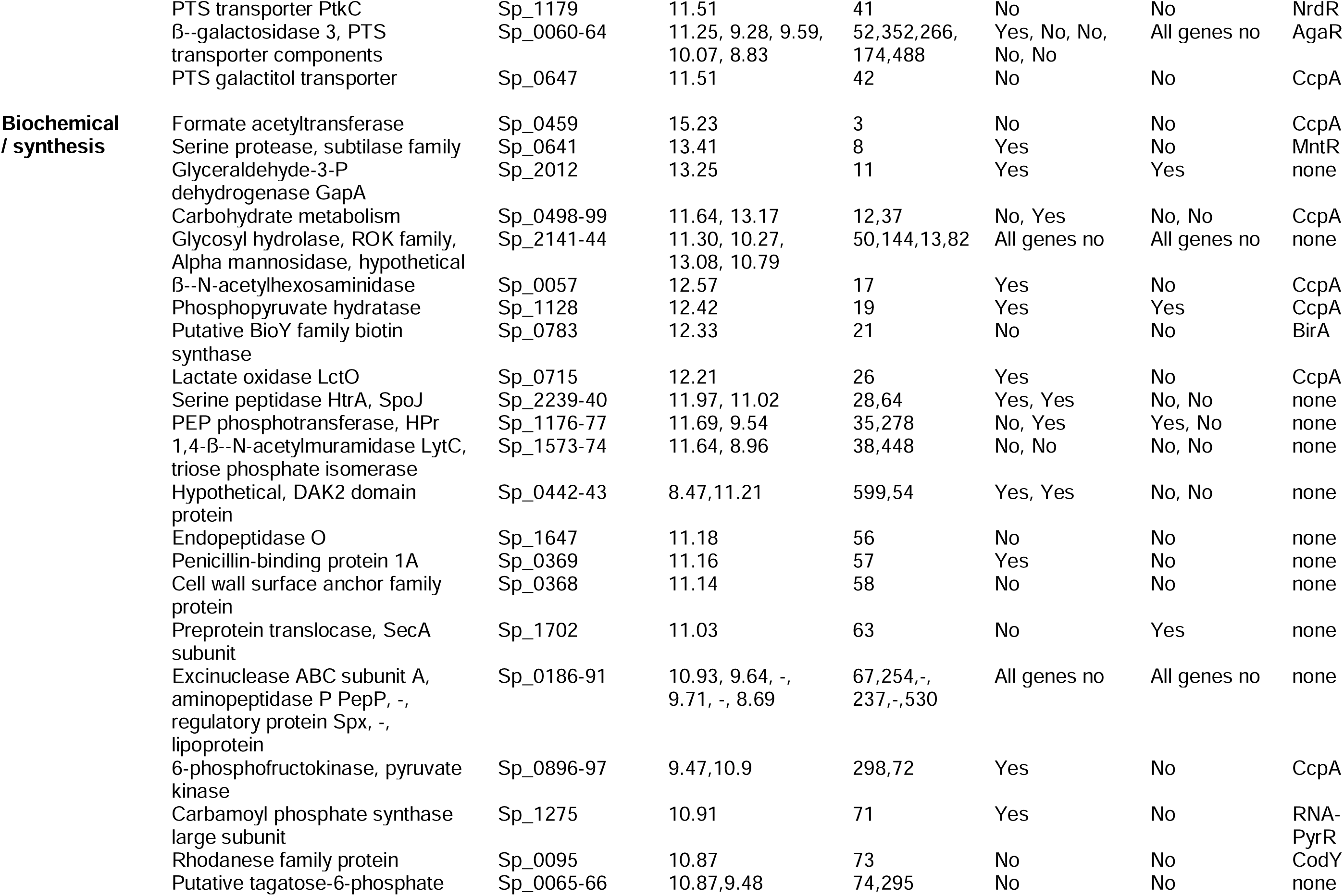
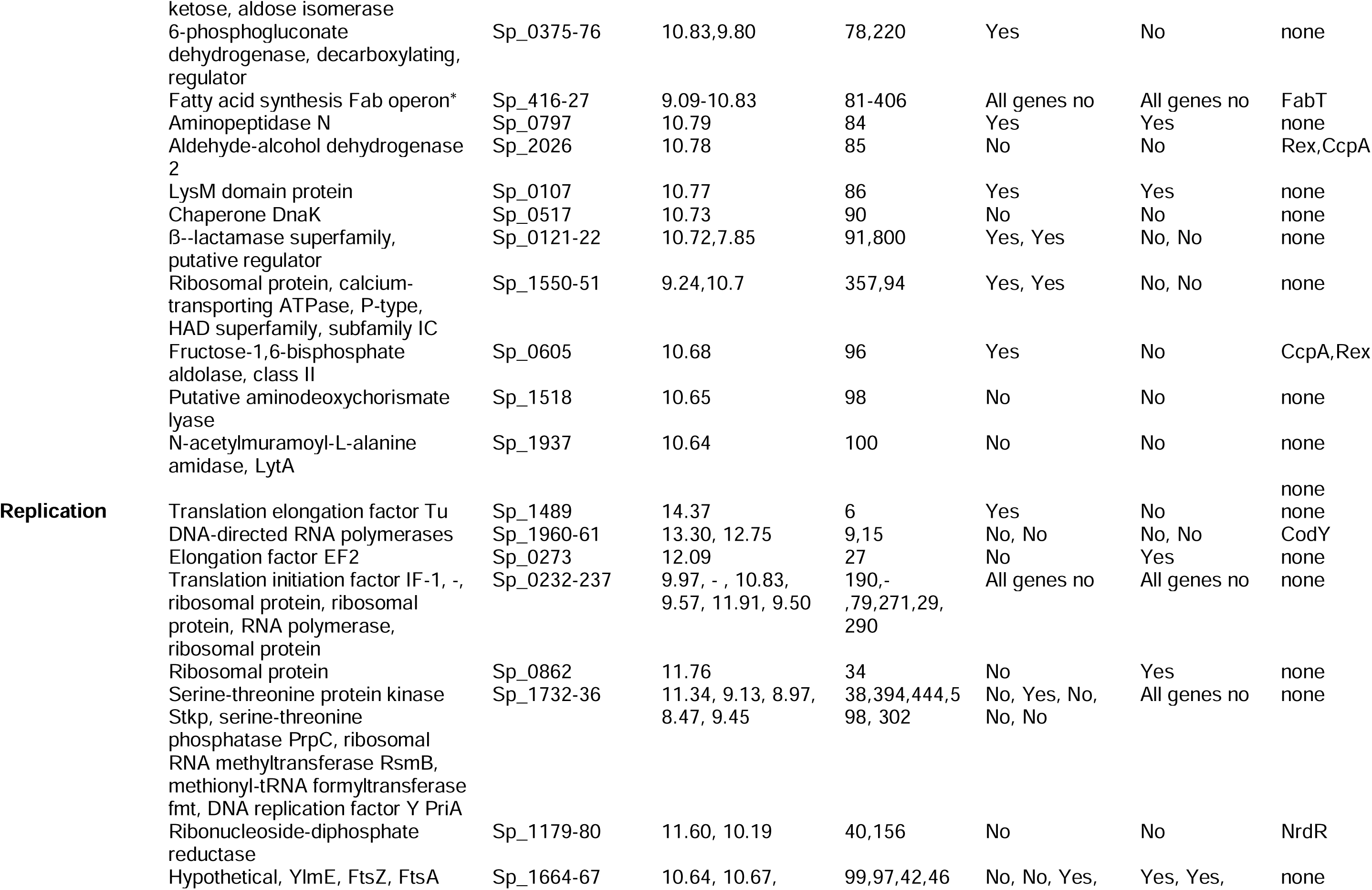
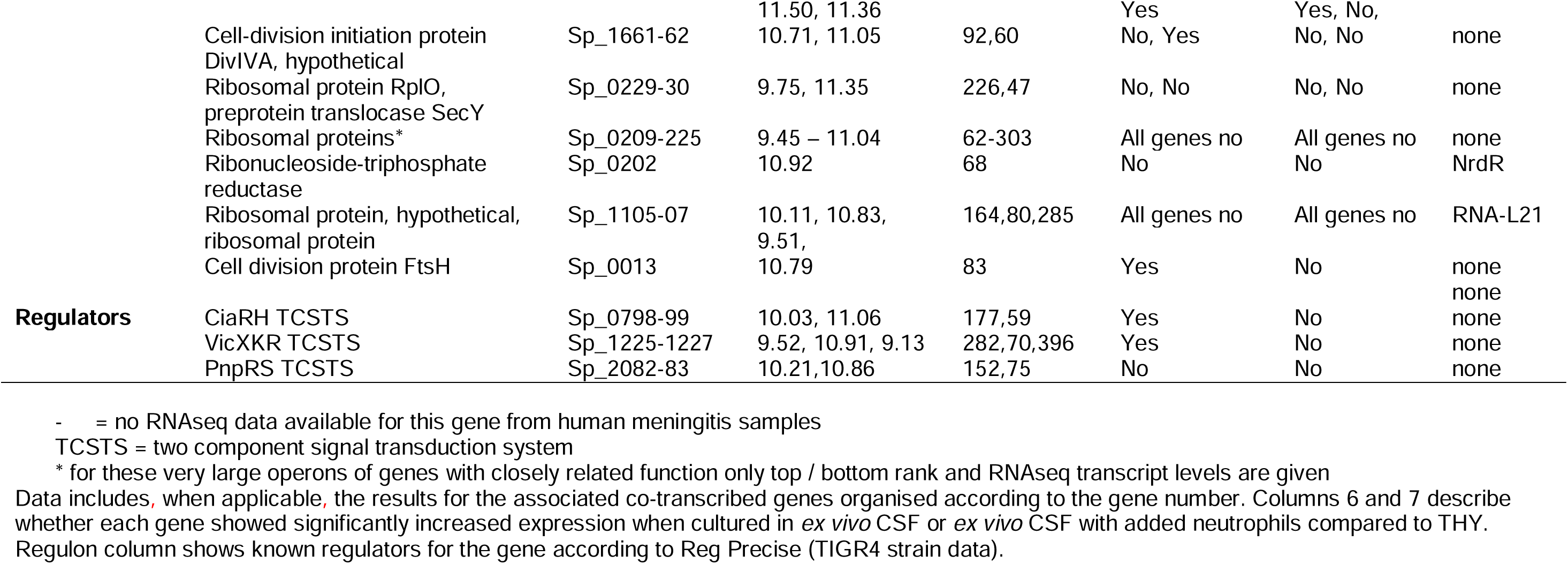
Highly expressed genes in CSF from humans with *S. pneumoniae* meningitis, identified as the 102 genes with RNAseq log_2_ transcripts 0.62 (1.5 SDs [3.73] greater than the mean [6.89] for the 2121 genes for which RNA was detected) divided into functional categories (by minant function for genes with multiple functions).

Classifying these highly expressed genes according to their annotated functions^28,29^ identified multiple genes encoding proteins involved in nucleic acid and protein biosynthesis and cell division (e.g. Sp_1489 and Sp_0273 translation elongation factors, Sp_1960 and Sp_1961 RNA polymerases, Sp_1966 and 1967 cell division proteins), consistent with active *S. pneumoniae* replication. Genes encoding known virulence factors were also highly expressed, including the choline binding proteins PspA (inhibits complement-mediated immunity)^30–32^ and PspC (involved in migration across the blood brain barrier and complement evasion),^12,33,34^ the manganese ABC transporter Psa (Sp_1648 – 1650, required for growth *in vivo* and resistance to oxidative stress),^35,36^ and neurotoxicity factors including pyruvate oxidase and pneumolysin^21,37–39^. In addition, multiple genes with no previously identified role in meningitis pathogenesis were highly expressed, including *bgaA* (encodes a ß-galactosidase), *lctO* (encodes a putative lactate oxidase), two component response regulators (eg *ciaRH, Sp_0661-62*), genes encoding the Ami and Sp_0090-92 ABC transporters, carbohydrate metabolism operons, and proteins with unknown or poorly described function (**Table 2**). Highly expressed genes during human meningitis rarely overlapped with genes identified by animal studies of meningitis or growth in CSF-mimicking media **(Supplementary Table 1)**; for example, none of the genes reported to be highly expressed during murine meningitis were found in the subset of 102 highly transcribed genes during human meningitis^40,41^. Further analysis using RegPrecise (https://regprecise.lbl.gov/)^42^ identified that highly transcribed genes in patient CSF belonged to regulons involved in manganese homeostasis, biosynthesis of fatty acids and deoxyribonucleotides, N-acetyl galactosamine utilisation, and ribosomal biogenesis (**Table 2, Supplementary Table 2**).

### Functional network analyses of the S. pneumoniae transcriptome during human meningitis

To provide further functional analyses of the RNAseq data, the top quartile of highly expressed genes in human meningitis were analysed using KEGG and STRING (https://string-db.org). KEGG network analysis showed upregulation of pathways involved in glycolysis, gluconeogenesis, RNA degradation, fatty acid biosynthesis, and pyruvate metabolic pathways (**Supplementary** Figure 1), likely to represent the most important metabolic pathways for pneumococcal response during meningitis. The STRING network analysis identified three functional co-expressed gene clusters representing gene networks involved in cell replication, metabolism, and genes known to be involved in meningitis pathogenesis (e.g. *ply*, *nanA* and *nanB*) (**Figure 1D**). The STRING network also identified two genes of unknown function, Sp_1802 and Sp_1804, outside of the three main networks, indicating their involvement in an additional mechanism of bacterial pathogenesis during meningitis.

### Identification of highly expressed genes during S. pneumoniae culture in CSF

Given the differences between *S. pneumoniae* transcriptomes in patients’ CSF versus *in vitro* ‘CSF-mimicking conditons’, we investigated whether highly expressed bacterial genes in patient CSF reflected adaptation to growth in CSF and/or interaction with phagocytes or more complex bacterial adaptations during meningitis pathogenesis. We compared the transcriptome of *S. pneumoniae* between culture in complete medium (THY) and *ex vivo* human CSF (from patients with normal pressure hydrocephalus), with or without addition of purified human neutrophils to partially represent the conditions found during pneumococcal meningitis^43^. The serotype 1 strain ST5316 (*GenBank* CABBZS000000000.1) was selected for these experiments as it was isolated from a human CSF sample, a close match to African meningitis strains^44–47^, and tractable for mutagenesis^48^. Principal component analysis showed wide separation of transcriptomes between the three *in vitro* conditions (**Figure 2A**), demonstrating a distinct bacterial transcriptional response for each condition. Differential gene expression analysis using DESeq2 identified 531 and 132 genes respectively with statistically increased expression in *ex vivo* CSF alone or *ex vivo* CSF plus neutrophils compared to THY, and 129 genes showing increasing expression in both *ex vivo* CSF and CSF plus neutrophils compared to THY (**Figure 2B and C**). Pathways analysis using KEGG of genes with increased expression in both *in vitro* CSF conditions compared to THY demonstrated over-representation of metabolic, biosynthesis of secondary metabolites, and purine metabolism pathways (**Figure 2D**), potentially representing pathways necessary for *S. pneumoniae* adaptation to CSF. Differential *S. pneumoniae* gene expression analysis using DESeq2 between culture in *ex vivo* CSF alone or in CSF plus neutrophils identified 466 pneumococcal genes with increased expression in the presence of neutrophils (**Figure 2D, supplementary data file 1**), representing (by KEGG analysis) enrichment for pathways including organisation of cell shape, replication, cytokinesis, and cell wall synthesis; these may represent specific *S. pneumoniae* adaptation to exposure to neutrophils.

**Figure 2:**
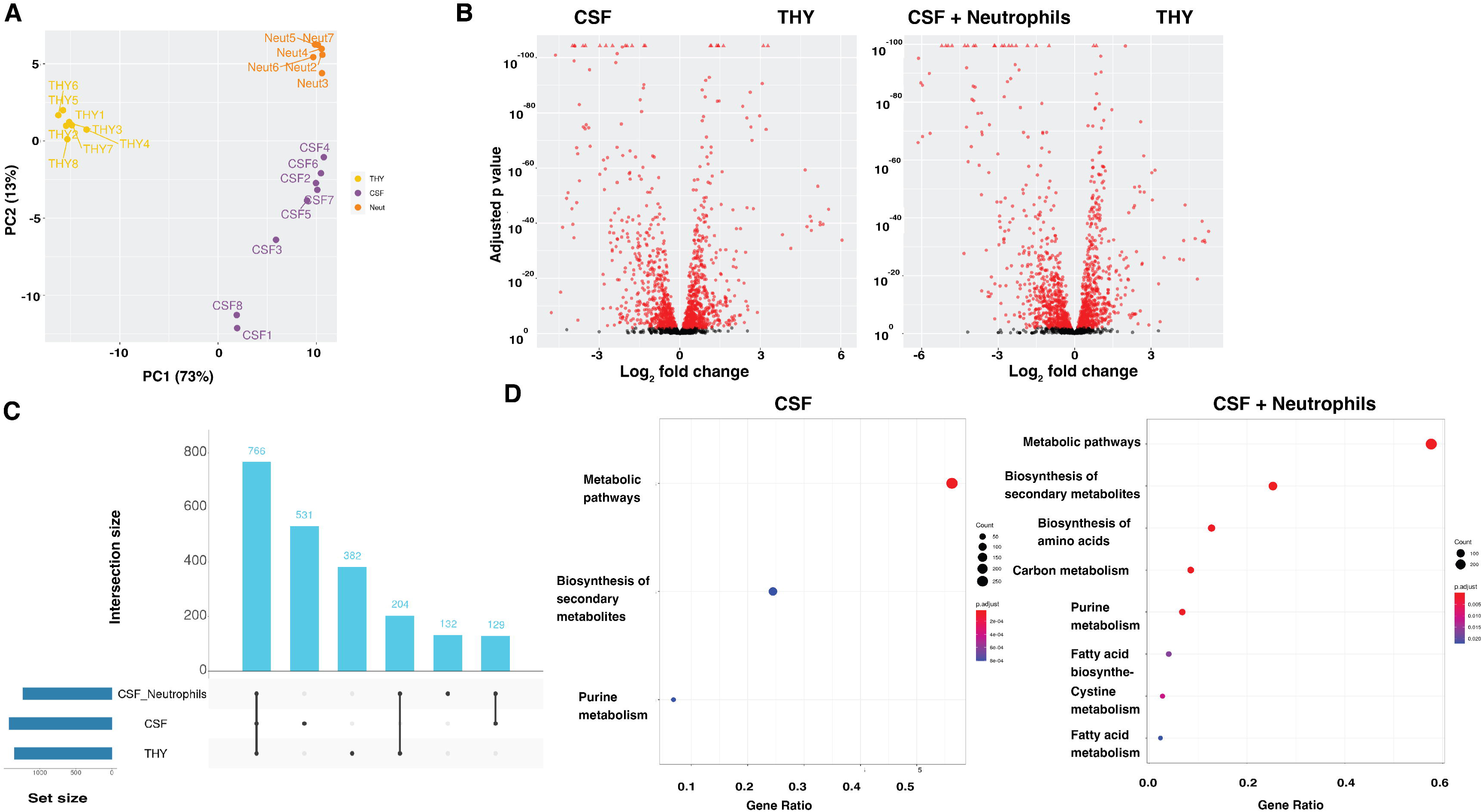
*S. pneumoniae* rapidly adapts to growth in CSF in vitro under growth-like conditions. (A) Principal component analysis of ST5216 transcription under 3 conditions: Todd-Hewitt broth (THY, yellow), human CSF (purple) or CSF with purified neutrophils at MOI 1 (orange) following 30 minutes of incubation at 37oC. (B) Volcano plot demonstrating the extent of differential gene expression between CSF and THY (Lt panel), and CSF + neutrophils and THY (Rt panel). Differentially expressed genes beyond the preset threshold shown in red. (C) UpSet plot quantifying numbers of co-expressed genes across the different conditions. All differentially expressed genes in the experiment were expressed at varying TPM in human CSF during meningitis. (D) Pathways analysis of differentially expressed genes in CSF conditions using KEGG. Dot size indicates number of pathway genes expressed (50-250), colour = adjusted p value.

Of the 102 highly expressed genes during human meningitis only 23 were significantly over-expressed in *ex vivo* CSF compared to THY (eg PspA, the CiaRH and VicXKR two-component sensor kinase systems, cell division proteins FtsH and FtsA, elongation factor Tu, and a range of metabolic proteins), 13 over-expressed in *ex vivo* CSF with added neutrophils (eg NADH oxidase and the Ami peptide ABC transporter), and 9 over-expressed in both *ex vivo* CSF conditions (eg Psa manganese uptake ABC transporter, GapA, and the hypothetical secreted protein Sp_1027) (**Table 2**). This shared requirement for high levels of expression of these genes during human meningitis and in *ex vivo* CSF conditions indicates they are involved in *S. pneumoniae* adaptation to growth in CSF and the presence of neutrophils. However, 57 genes highly expressed during human meningitis were not differentially expressed under *ex vivo* CSF conditions, including genes encoding Ply, PspC, and SpxB, along with ß-galactosidase (BgaA), ZmpB, fatty acid biosynthesis enzymes, the PnpRS two-component sensor kinase system, and proteins of unknown function. These genes potentially represent bacterial factors required for CNS infection that are harder to identify and characterise *in vitro*, without obtaining ex vivo bacterial RNAseq data from infected human CSF, and hence have not been previously characterised through animal or *in vitro* model systems.

### Selection and mutation of two genetic loci highly expressed during human meningitis

To further investigate genes that were highly expressed in human disease but not in *in vitro* meningitis-like conditions, *bgaA* and the operon *Sp_1801-05* were selected. Proteins encoded by these genes have no previously described role during meningitis, but were among the most highly expressed genes in human meningitis samples, and were also co-expressed in gene networks identified by the STRING analysis, *bgaA* within the metabolic cluster and Sp_1802 and Sp_1804 separate from the main network clusters (**Figure 1D**). *bgaA* was the second most highly expressed gene in the human meningitis dataset (median log_2_ TPM 15.25), and encodes a ß-galactosidase that aids *S. pneumoniae* growth in semi-defined media, adherence to bronchial epithelium, and inhibits complement activity^49^. Sp_1801-05 is a five gene operon with a mean log_2_ TPM of 10.94 (range 9.18 to 12.56), and is conserved across Gram positive species. The function of proteins encoded by Sp_1801-05 is poorly understood, with published data only available for Sp_1804 (described as encoding a haemin binding protein)^24,50–52^. *In silico* analysis using multiple proteomic mapping tools (**Supplementary Table 3, Supplementary** Figure 2) suggested Sp_1801-05 proteins are involved in stress response; Sp_1802 and Sp_1804 are related to alkaline stress response proteins, and Sp_1805 has similarity to CsbD a protein involved in bacterial resistance to environmental stresses^53^.

### *In vitro* characterisation of Δ*bgaA* and Δ*Sp_1801-05* strains

Serotype 1 *S. pneumoniae* ST5316 mutant strains containing deletions of *bgaA* or the *Sp_1801-05* operon were constructed recently described techniques to transform serotype 1 strains (**Figure 3A**).^48^, and their phenotypes investigated in assays relevant for meningitis pathogenesis. Both Δ*bgaA* and Δ*Sp_1801-05* strains grew at a similar rate to the wild-type (WT) strain in THY and under conditions of osmotic and cation stress (**Figure 3B**). Compatible with potential roles suggested by *in silico* analysis for Sp_1802 and 1804 in responding to pH stress, the Δ*Sp_1801-05* strains had slightly delayed growth in THY media under high (8.0) and low (6.8) pH conditions (**Figure 3B**). Both the Δ*bgaA* and Δ*Sp_1801-05* strains had significantly delayed growth in *ex vivo* CSF compared to the wild-type strain (**Figure 3B**). Compatible with previously published data^49,54^ the Δ*bgaA* strain was more sensitive to opsonisation with complement C3b/iC3b in serum. CSF contained too little complement to further assess opsonisation in this compartment (**Figure 3C**). In addition, the Δ*Sp_1801-05* strain was also more sensitive to opsonisation with complement (**Figure 3G and H)**. A monolayer model of Human Brain Microvascular Endothelial Cells (HBMEC) and a multi-cellular transwell model of the blood brain barrier (BBB) including HBMEC, pericytes, neurons and microglia (**Supplementary** Figure 3) were used to evaluate if either protein was involved in transmigration or disruption of the BBB. When measured by electrical impedance neither mutant strain showed differences in the early disruption HBMEC monolayer tight junctions caused by *S. pneumoniae*. HBMEC cell death was marginally delayed after infection with the Δ*bgaA* and Δ*Sp_1801-05* strains compared to WT control, but remained greater than the Δ*ply* control (**Figure 3D**). Electrical impedance was maintained across the multi-cellular transwell BBB model by both mutant strains compared to WT and the Δ*ply* control (**Figure 3E**), but all strains showed similar levels of transmigration across the BBB layer (**Figure 3F**).

**Figure 3:**
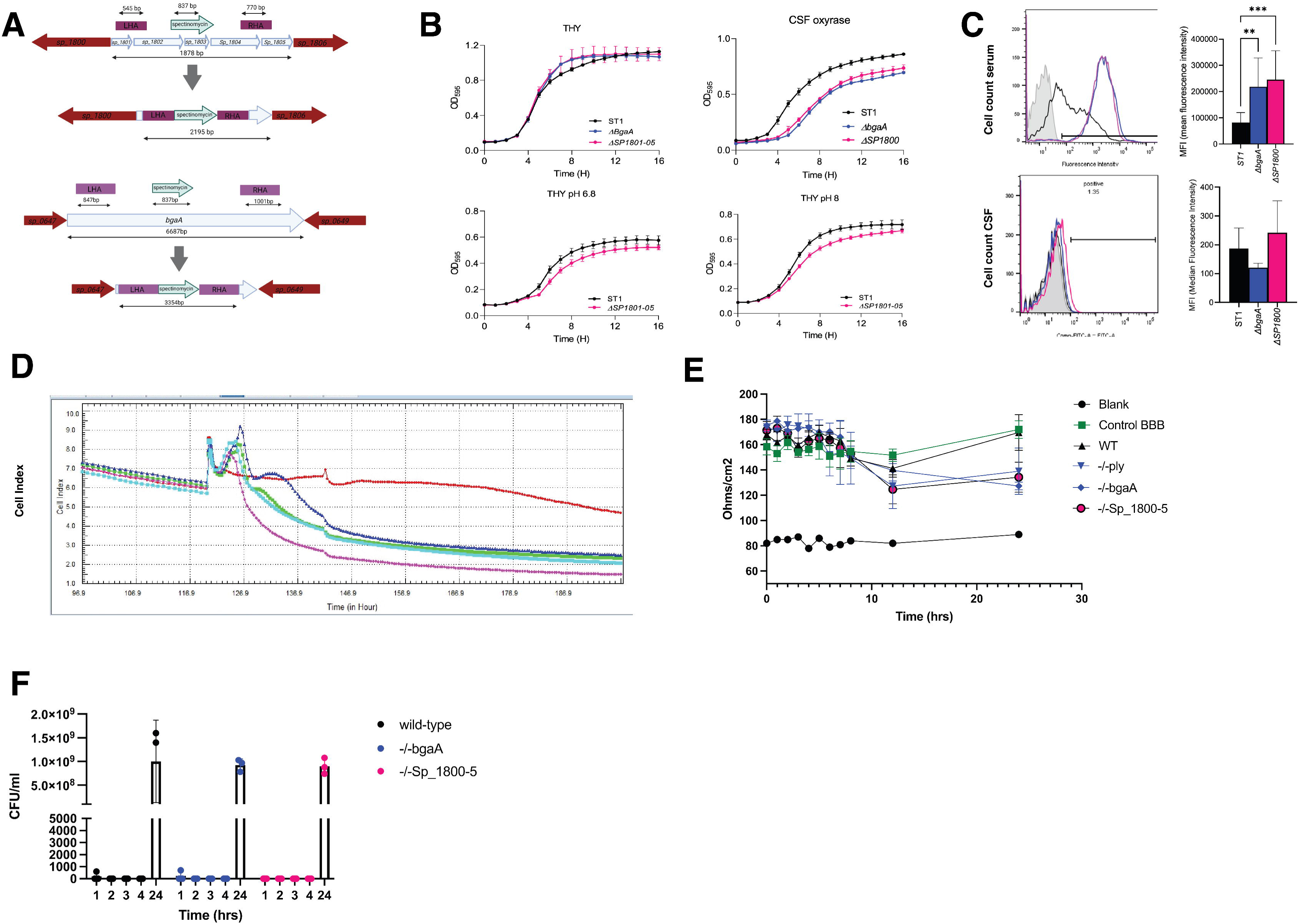
Construction and phenotyping of bgaA and Sp_1800-5 gene deletion mutants in *S. pneumoniae* serotype one. (A) Gene-deletion mutant construction of ΔSp_1800-5 and ΔbgaA in *S. pneumoniae* serotype 1 strain 519/43 316 through insertion of a spectinomycin inactivation cassette. (B) Growth of ST1, compared to ΔbgaA and ΔSp_1800-5 in Todd-Hewitt media, CSF supplemented with oxyrase, or THY in acidic (pH 6.8) and alkaline (pH 8) conditions. Optical density at 520 nm recorded hourly to 18 hours. (C) Complement binding of ST1 compared to gene deleted mutations in serum and CSF, measured flow cytometry. Left panels proportion of FITC-labelled cells (C3 bound) m right panels comparison of MFI between conditions. Top row serum, bottom row, CSF. (D) Kinetics of *S. pneumoniae* disruption of endothelial tight junctions in a monolayer of HBMEC cells measured in the XCELligence system, using cell index (y axis) as a measure of electrical conduction across cells. Control hBEMC cells in red, WT (purple), Δply (dark blue), ΔbgaA (green), ΔSp_1800-5 (light blue). (E) Electrical impedance (Ohms/cm2, y axis) across a 4 cell-type *in vitro* transwell model of the BBB over time between WT and gene deletion mutations, including Δply, compared to uninfected cells and blank transwell to 24 hours. (F) Bacterial growth (CFU/ml) in the collecting chamber of the BBB transwell model over time.

### Virulence and BBB transmigration are attenuated in both ΔbgaA and ΔSp_1801-05 mutant strains

The effects of mutation of *bgaA* or *Sp_1801-05* was assessed in a recently developed mouse model of brain infection involving nasopharyngeal translocation of *S. pneumoniae* to the brain^55^. Both the Δ*bgaA* and Δ*Sp_1801-05* strains successfully colonised the nasopharynx and were able to reach the olfactory epithelium and olfactory bulb with similar CFU levels at all three sites compared to the WT strain (**Figure 4A**). However, although the occasional mouse infected with the mutant strains had detectable CFU at earlier timepoints, by ten days post infection, no mouse infected with either mutant strain had detectable CFU in their brain tissue (**Figure 4A**). In contrast, brain infection occurred in 50% of mice infected with WT, demonstrating that both *bgaA* and *Sp_1801-05* were required for brain infection. Next, we tested the virulence of the two mutant strains in a zebra fish meningitis model requiring direct injection of *S. pneumoniae* into the hindbrain CSF ^55,56^. Zebra fish injected with the Δ*bgaA* strain had improved survival compared to those injected with the Δ*Sp_1801-05* or WT strains, although there were no differences in either bacterial CFU recovered from the fish brain or neutrophil ingress (**Figures 4B and C**).

**Figure 4:**
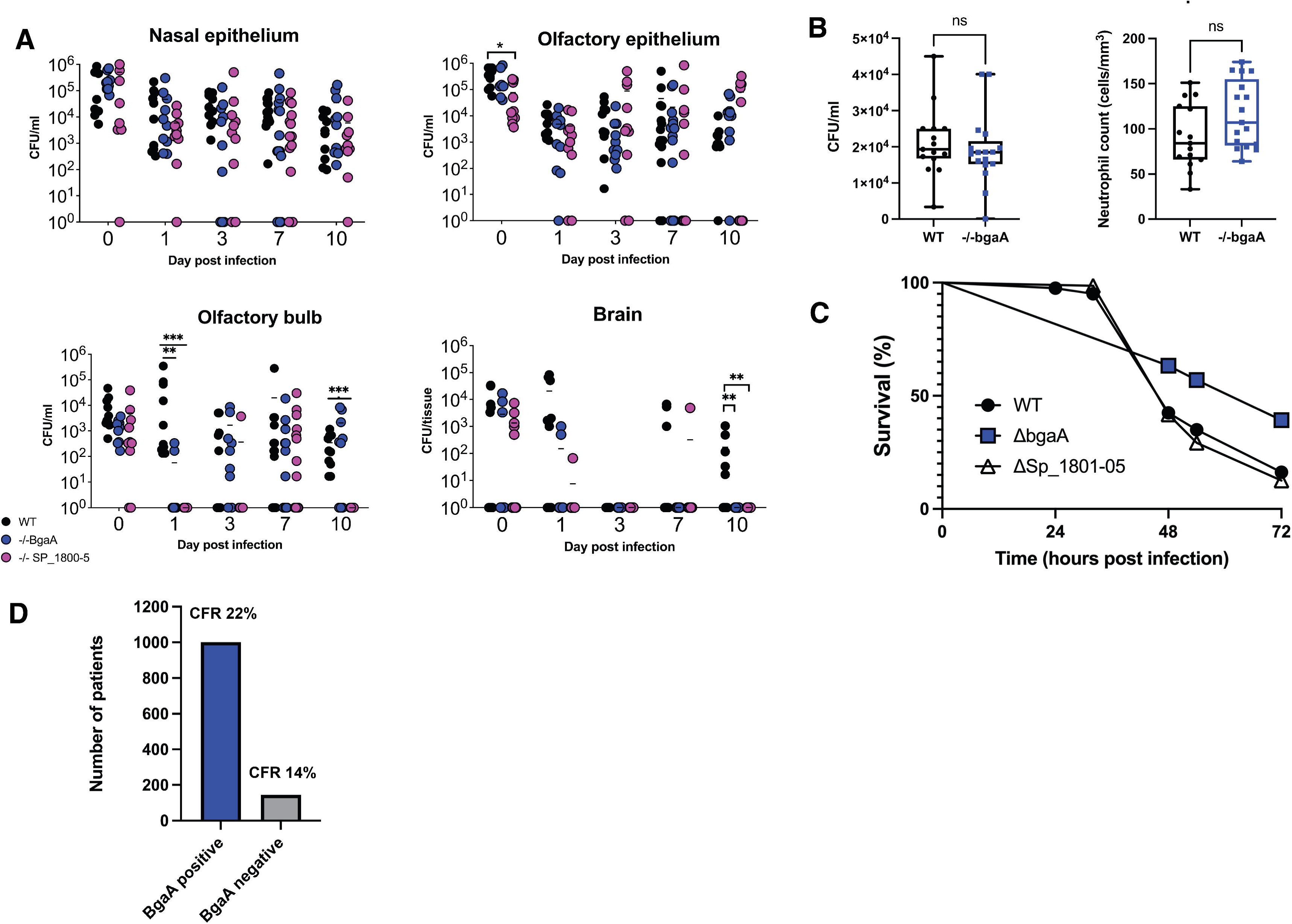
*bgaA* and Sp_1800-5 are required for bacterial survival in the CNS and are important for virulence. (A) Transmigration of wild-type, ΔbgaA (blue) and ΔSP_1800-5 (pink) from nasal epithelium, olfactory epithelium and bulb, and brain in a murine trans-nasal meningitis model. Bacteria were quantified at days 1,3,7 and 10 post inoculation by colony counting in each compartment. (B) Bacterial counts (CFU/ml, Lt panel) and neutrophil ingress (cell counts by con-focal microscopy) to Zebrafish hind brain 5 hours following inoculation with WT (black) compared to Δ*bgaA* (blue). (C) Survival of zebrafish following hindbrain inoculation with WT, ΔbgaA and ΔSp_1800-5. (D) Numbers of patients in the Dutch meningitis database with sequence-confirmed *S. pneumoniae* containing BgaA (blue) and without BgaA (grey). Case fatality rate (CFR) for each strain is given in the figure.

### The presence of bgaA was associated with poorer outcome in PM human patients

To test for association between infection with strains containing *bgaA* or Sp_1801-05 and clinical outcome, genome sequences for 1144 *S. pneumoniae* strains recovered from Dutch meningitis patients were analysed. The Sp_1801-05 operon was present in 702 (61.4%) genomes, but this was not associated with an increased mortality (22% versus 20%, P = 0.682)). *bgaA* was present in 1000/1144 (87.9%) *S. pneumoniae* strains across all serotypes, and infection with *bgaA* positive strains was associated with a lower CSF white cell count (2653 cells/mm^3^ range 545-7849 versus 4330 cells/mm^3^ range 975-9725), P = 0.037). In patients where the mortality was known (n= 1035), infection with strains containing *bgaA* was associated with a higher mortality (204/909, 22%) compared to *bgaA* negative strains (17/126, 14%), Fisher exact test p = 0.008) (**Figure 4D**, **Supplementary Table 4**). These data further support that *bgaA* has an important role during the pathogenesis of *S. pneumoniae* meningitis.

## Discussion

Transcriptomic approaches offer a comprehensive insight into bacterial processes during infection, potentially shedding light on factors crucial for virulence. To our knowledge there are no published studies on the overall *S. pneumoniae* gene transcriptome during invasive human infection. Our *S. pneumoniae* RNAseq of CSF obtained from patients with pneumococcal meningitis represents a distinctive dataset, revealing several new findings including: (i) a remarkable consistency in highly expressed *S. pneumoniae* genes among patients, indicating activity in the metabolic pathways likely vital for bacterial survival and replication during meningitis; (ii) many highly expressed genes in meningitis were not highly expressed during culture in *ex vivo* human CSF or *in vitro* CSF-mimicking conditions, underscoring the need to use human samples for their identification; (iii) mutant phenotype analysis demonstrating two human meningitis-specific highly expressed genetic loci, *bgaA* and *Sp1801-05*, were required for brain infection in a mouse model; and (iv) genome analyses indicated *bgaA-*containing strains were associated with increased severity in human meningitis. Overall, RNAseq of huamn CSF samples has identified the metabolic pathways that are active during *S. pneumoniae* meningitis and previously unsuspected important roles for two genetic loci during brain infection.

The importance of the genes showing increased expression only during human meningitis for disease pathogenesis was supported by the phenotype data obtained for *bgaA* and Sp1801-05 operon gene-deletion mutant strains. *bgaA* (the second most highly expressed gene in human meningitis) encodes BgaA, one of several *S. pneumoniae* exoglucosidases and which cleaves N-terminal galactoses linked to glucose or N-acetylyglucosaminidase on host glycoproteins. BgaA has roles in bacterial metabolism, cell adherence and avoidance of opsonophagocytosis^49,57^. There are almost no data on the function(s) of the Sp1801-05 operon, but our *in silico* analysis suggests it is involved in the bacterial response to environmental stress, and this is supported by our data showing the Δ*Sp_1801-05* strain has delayed growth under lower or higher pH conditions.^50^ In addition, the D39 homolog of Sp_1804 (SPD_1590) may play a role in iron transport and adherence to human lung epithelial cells.^50^ Using a recently developed murine model of brain infection that replicates *S. pneumoniae* spread to the brain via the cribriform plate, we showed that both Δ*bgaA* and Δ*Sp_1801-05* strains failed to establish sustained infection of the brain supporting the hypothesis that they have a significant role during meningitis pathogenesis. Phenotype analyses indicated both the Δ*bgaA* and Δ*Sp_1801-05* strains had delayed growth in the CSF and increased sensitivity to opsonisation with complement compared to the wild-type strain. Both phenotypes are likely to be important for the pathogenesis of meningitis; rapid growth in CSF will be necessary for *S. pneumoniae* to establish infection, complement proteins are essential mediators of innate immunity to *S. pneumoniae*^58^ and are present in high concentrations in CSF during meningitis^59^. The role of *bgaA* for growth in CSF is surprising,

as glucose is the dominant carbon source in CSF rather than N-linked glycans which are the substrate for *bgaA*^49^. The effect of *bgaA* on complement resistance has been described previously^60^ and is thought to be mediated by de-glycosylation of complement proteins. The pathogenic role of bgaA in pneumococcal meningitis. The impaired growth of the Δ*Sp_1801-05* strain in CSF would be compatible a role in stress response and/or low iron levels in CSF. How Sp_1801-05 can affect complement activity is not clear and requires further investigation. Importantly, meningitis caused by *bga* containing strains was associated with reduced CSF white cell concentrations and higher mortality further supporting an important role for *bga* during meningitis pathogenesis. These data were not controlled for differences in distribution of *bgaA* between serotypes, but *bgaA* is widely distributed amongst pneumococcal serotypes and was absent in a minority of strains making it less likely that the association with mortality is caused by background genotype independent of *bga*.

CSF provided a sample directly from the site of infection containing a high bacterial load due to disease severity and lack of prior antibiotic use, and this allowed successfully isolation of bacterial RNA from a proportion of meningitis patients. For practical and ethical reasons repeat lumbar punctures to identify temporal changes in transcriptome response were not possible. The data analysis faced technical challenges. Our patients were infected with different *S. pneumoniae* serotypes, providing challenges to mapping and annotation of pneumococcal genes. Most of our subjects were HIV positive, and we are unable to determine if immunosuprresion through HIV infection affected which *S. pneumoniae* genes were highly expressed in CSF.

We chose to map all samples to a single geographically relevant serotype, and transformed gene names into the TIGR4 nomenclature to facilitate functional annotation but were limited by the lack of an agreed pan-serotype functional annotation system for *S. pneumoniae*. Furthermore, the significant methodological differences in obtaining human CSF RNAseq data compared to from *in vitro* cultures precluded conventional statistical comparison of the datasets. Instead, we have used SD 1.5+ above the median TPM for all genes across all samples to identify 102 highly expressed genes during human meningitis, or for the STRING and KEGG analyses the top quartile of expressed genes to increase the potential range of networks identified. Generally, when highly expressed genes are part of an operon, the rest of the operon also showed higher expression levels (eg the *fab*, *Sp_0090-92*, *ami*, *Sp_2141-44* operons) indicating the data reflect biologically relevant upregulation of gene expression.

Despite the inherent variation caused by different infecting strains, host background, and variable timing of presentation there was significant consistency in which genes were highly expressed across subjects. These included genes encoding proteins with known roles in meningitis pathogenesis (eg *ply*, *pspA*, *pspC*), but also many genes with no known role. The pathogenesis of meningitis involves *S. pneumoniae* growth in CSF containing a large influx of neutrophils, conditions which we replicated using culture in *ex vivo* CSF,^61^ brain damage in meningitis is due to both direct pathogen-mediated, and secondary damage from neutrophil-mediated inflammation.^61–63^ Our *in vitro* conditions identified a large number of genes that were differentially expressed compared to *S. pneumoniae* culture in THY, which are likely to represent metabolic adaptation to CSF and/or response to interactions with neutrophils that will be investigated in the future. The large number of highly expressed genes only identified in the meningitis dataset emphasises the importance of obtaining data from disease subjects to fully understand pathogenesis. These genes could reflect the greater complexity of *S. pneumoniae* / host interactions during actual meningitis compared to *ex vivo* CSF, as well as differences due to evolving gene expression during human disease compared to short term *ex vivo* culture.

To conclude, we have used direct RNAseq from clinical samples to identify the *S. pneumoniae* genes and gene networks that are highly expressed during human meningitis. Comparison of the data to *in vitro* transcriptomic studies in *ex vivo* CSF identified multiple genes that were specifically highly expressed during human meningitis, including *bgaA* and the Sp_1801-05 operon. Subsequent mutational analysis demonstrated that *bgaA* and Sp_1801-05 are important for establishing brain infection in a mouse nasopharyngeal to meninges translocation model. Our work thus provides a road-map for identifying important novel mechanisms required for the pathogenesis of *S. pneumoniae* meningitis, data needed to help develop future therapeutic interventions against this devastating disease.

## Methods

### Participants

Adults and adolescents presenting to Queen Elizabeth Central Hospital in Blantyre, Malawi with subsequently proven bacterial meningitis caused by *S. pneumoniae* between 2011-2013 were included (Current Controlled Trials registration ISRCTN96218197)^25^. All CSF and blood samples were collected at the bedside prior to administration of parenteral ceftriaxone 2g BD for 10 days.^7,64^

### Ethics

All participants or nominated guardians gave written informed consent for inclusion. Ethical approval for the transcriptomics study was granted by both the College of Medicine Research and Ethics Committee (COMREC), University of Malawi, (P.01/10/980, January 2011), and the Liverpool School of Tropical Medicine Research Ethics Committee, UK (P10.70, November 2010) Committee, Liverpool, UK.

### Procedures

Routine CSF microscopy, cell count, and CSF culture was done at the Malawi-Liverpool-Wellcome Trust Clinical Research Programme laboratory in Blantyre, Malawi as previously described^25^. Culture negatives samples were screened using the multiplex real-time polymerase chain reaction for *S. pneumoniae*, *N. meningitidis* and *Haemophilus influenzae type b* (Hib) kit from Fast-Track Diagnostics (FTD Luxemburg) according to the manufacturer’s instructions, bacterial load estimated from Ct values, using standards previously generated.^65^ We collected 2.5 ml of CSF and whole blood for transcriptional profiling in blood PAX-gene® (Pre-AnalytiX, Qiagen, USA) tubes, incubated for 4 hours at room temperature following the manufacturers instructions, and stored at –80 degrees Celsius. In-hospital HIV testing was done on all patients by the clinical teams using point-of care Genie™ HIV1&2 test kits (BioRad, USA).

RNA was extracted from human CSF samples using the PAXgene® Blood miRNA kit (Pre-Analytix, Qiagen, USA) according to the manufacturer’s instructions, with an additional mechanical disruption step of CSF samples to disrupt the pneumococcal cell wall at 6200 rpm for 45 seconds in the Precellys evolution tissue homogenizer (Bertin Instruments). The extracted RNA was quantified and RNA Integrity Number (RIN) scores calculated using RNA Tapestation 4200® (Agilent, USA) and Nanodrop® (Thermoscientific, USA). CSF samples were selected for additional ribodepletion and bacterial RNA sequencing where a bacterial 16S spike was seen on the Tapestation trace, irrespective of overall RIN. Ribodepletion was done with the Illumina RiboZero Gold kit, following the manufacturers instructions.

Extracted RNA samples underwent library preparation for total RNA sequencing in samples where the pre-ribodepletion RNA was >1ng/1ul using with Kapa RNA hyperPrep kit (Roche), followed by 75 cycles of Next-generation sequencing with NextSeq® (Illumina, USA) by the Pathogen Genomics Laboratory at University College London.

### Bacterial mapping & annotation

The paired-end libraries were aligned individually onto 70 *S. pneumoniae* genomes (NCBI, 8 April 2019); these genomes were described as complete on the date of access. Alignment was performed by RNA-STAR (v2.6.0a)^1^ with the following options: (i) alignIntronMax 1 and (ii) sjdbOverhang 40. Alignment rate (i.e. fraction aligned to individual genome) was calculated for every genome per sample. For every sample, a cut off was defined to separate genomes with high alignment rate and non-high rate using Kernel density distribution in R (R v3.5.2). Complete alignment and alignment rate are listed in **Supplemental data 1**. For the selected genomes, the aligned reads were then summarized (featureCount v1.6.3) according to the chimeric annotation file in stranded, multimapping (-M), fractionized (--fraction) and overlapping (-O) modes^2^. Additionally, we listed homologous genes among the pneumococcal genomes including GCF_003003495 (*S. pneumoniae* D39V) by Mauve v20150226^3^; homologous genes were defined as having common coverage at least 60% and identity at least 70%.

In situations where comparison with publicly available data was performed, the data was mapped to the *S. pneumoniae* TIGR4 reference genome (NC_018630) using nf-core/rnaseq v3.4.

Gene expression was normalized against gene length and library size, or as TPM (transcript per million). For samples with more than one high alignment rate genomes, we included only homologous genes shared among the genomes. Moreover, the recently completed *S. pneumoniae* D39V annotation^4^ was used as the base of analysis by using the homologous genes between the selected genome(s) and D39V. To group pneumococcal genes into highly expressed genes and non-highly expressed genes (i.e.: the high cut-off), we used k-means clustering to the normalized gene expression to divide genes into twelve clusters^5^. From the twelve clusters, high clusters were selected so that the high genes were at least formed the top 8% highly expressed genes. For samples 183C and 283C, high and non-high genes were grouped by Kernel density distribution. Enrichment test were performed by the built-in function. Corresponding *p*-value of the enrichment test were adjusted by Bonferroni correction.

### Generation of gene-deletion mutants

Gene deletion mutants of *S. pneumoniae* serotype 1 were generated as previously described.^48^ In brief, a serotype 1 strain 519/43 ST5316, isolated in 1943 from a patient CSF in Denmark, acquired from the Statens Serum Institute was used. ^48^ A spectinomycin cassette was inserted into the gene region inactivate either BgaA or the operon *Sp_1801-5* usingprimers BgaA FW1: 5’ tttgcggccgcggccattggaatcggaaaagagtttata 3’ Bga1RV: 5’ tgtccatgcagtcaataaacagccaaggatccacttactttctcataaaccagttgctgcgg 3’, BgaAFW2: 5’ ttggctgtttattgactgcatggacaggatccacttactttctcataaaccagttgctgcgg 3’ BgaRV2: 5’ ccttccagtacttgttcgcctcgttgcggccgcaaa 3’, _LHAFW: 5’tttgcggccgctgtattcgctggtttagtcggggcat3’, LHARV: 5’ cggcaatcgaaggttttgttagaagtttggatcctactgttgtagctcacgaaatcaaagggga 3’, and RHAFW 5’ cggcaatcgaaggttttgttagaagtttggatcctactgttgtagctcacgaaatcaaagggga 3’, RHARV: 5’ cgctgtagaaggtgctgtagaaggtgttaaaagcggccgcaaa3’ respectively. Correct introduction in the chromosome was confirmed by using primers BgaSCN1: actaggttgtcataccatgtataccacttg and BgaSCN2: actattttgttccagactctttatcttctattt for *bgaA* and primers upSCN1: 5’ ttattgctggaggtcttattggtctcttgg3’ and downSCN2: 5’ gcaattgggaatctctagctttttgttttctgag3’ for the modified region, sp_1801-1805. All positive clones for insertion were confirmed by sequencing.

### Bacterial strains and growth conditions

*Streptococcus pneumoniae* was cultivated in Todd-Hewitt broth (Roche), supplemented with 0.5% yeast extract (THY) at 37°C in 5% CO_2_ to optical density (OD) of 0.5 at 620 nm. Genetically-modified pneumolysin-deleted mutant bacteria were selected from growth on Colombia agar (Oxoid, UK) supplemented with 5% horse blood and 100 µµg/ml of Spectinomycin as previously described^48^, grown in THY under spectinomycin selection to OD 0.5. Bacterial stocks were enumerated by plating serial dilutions on blood agar and stored in 80% glycerol at –80°C.

### S. pneumoniae growth in human CSF

Human CSF samples were a kind gift to ECW from DvdB at the Amsterdam Medical Centre, University of Amsterdam, The Netherlands. Surplus normal lumbar CSF was obtained from diagnostic lumbar punctures for patients with a clinical diagnosis of normal pressure hydrocephalus or benign intracranial hypertension with consent, snap frozen, shipped at – 80°C and thawed on ice to preserve active complement. Complement was depleted from serum and CSF where required by heating to 65°C for ten minutes. For all growth experiments, bacteria were thawed from stocks, washed twice in PBS and re-suspended at an OD of 0.1. Growth in THY was used as a positive control, five technical replicates were undertaken for all conditions. Growth was measured using a Tecan Spark plate reader (Tecan, USA) at 37°C in 5% CO_2_ with shaking at 200 rpm for 24 hours. Optical density readings at 620 nm were taken at 30-minute intervals. Samples were serially diluted and colony forming units were plated on blood agar in parallel every 2 hours for the first 8 hours of culture using adapted Miles & Misra method.^66^

Fresh human neutrophils were extracted from whole blood of healthy lab donors by negative selection using the MACSxpress® system (Miltenyibiotec, USA) according to the manufacturer’s instructions. Erythrocytes were depleted post neutrophil isolation by incubation for 8 minutes with 1X Invitrogen RBC lysis buffer (ThermoFisher, USA) prior to all experiments. Neutrophil viability was assessed by Trypan blue staining. Neutrophils were counted using a cell chamber and adjusted in all experiments to 2×10^6^ cells/ml. Neutrophils were re-suspended in HBSS with 10% serum or CSF and kept at 37°C until use (<4 hrs). All experiments used an MOI of 1.

### RNA extraction and sequencing of S. pneumoniae in CSF culture

Bacteria were cultured in THY until mid-log phase, pelleted at 4000g for 5 minutes, washed in PBS three times and resuspended in 1ml of either fresh THY, CSF warmed to 37°C, or CSF with 1×10^6^ fresh neutrophils. Eight replicates of each condition were used. All samples were incubated for a further 30 minutes in 5% CO_2_ at 37°C, before being incubated directly into 2 mls of RNA*later to* preserve bacterial RNA. All samples were incubated for 4 hours in RNA*later* at room temperature and then frozen at –80°C. *S. pneumoniae* RNA was extracted following a method developed by Mann et al, using the MirVana phenol based extraction kit (Thermofisher) as previous reported.^67^ Briefly, samples were thawed, pelleted and RNA protection media was removed. Cell lysis buffer was applied, samples were placed in a FastPrep MatrixE tube, undergoing mechanical cell wall disruption in a Precellys machine speed of 6200 rpm for 45 seconds. Homogenates were incubated in a water bath for 10 minutes at 70°C, cooled on ice and then pelleted at 12k x g, 5 min, 4°C in pre-cooled microfuge. The supernatant containing RNA was removed and passed through a Qiashredder (Qiagen) for 2 minutes at 12k x g, in the same 4°C in pre-cooled microfuge.

RNA extraction was completed using the MirVana kit (ThermoFisher) following the manufacturers instructions. Following nucleic acid extraction, TurboDNAase enzyme and buffer (1:10 ratio) (ThermoFisher) were added to each sample and incubated at 37°C for 30 minutes. RNA quality was quantified using the Tapestation/BioAnalyser. Ribosomal RNA was depleted using the bacteria rRNA depletion kit (New England Biolabs), with the addition of human rRNA depletion beads for the samples containing *S. pneumoniae* cultured in CSF + Neutrophils. Libraries were prepared as previously using the Kapa kit for total RNA and sequenced by the Pathogen Genomics Unit (PGU) at University College London.

### Bioinformatics & data analysis

RNA-seq reads were using nf-core/rnaseq v3.4. Alignment was performed using HISAT2 after read trimming and quality control steps using fastp. The nf-core/rnaseq pipeline implements best practices for standardized RNA-seq analysis using Nextflow. The pneumococcal transcriptome was compared under different *in vitro* conditions using DESeq2.

*In vitro* laboratory data were visualised using GraphPad Prism version 9, data were summarised using medians and range, different conditions were compared using pair-wise comparisons Mann-Whitney-U tests.

### Data availability

Summaries of the sequenced, mapped data and analysis for both the *in vivo* human pneumococcal transcriptome from meningitis patients and *in vitro* transcriptome (**Supplemental data 1**) are available from the UCL data repository rdr.ucl.ac.uk URL DOI: https://doi.org/10.5522/04/25721628.v1

## Meningitis models

### In vitro blood brain barrier models

#### Ethical approval

Human foetal brain tissues from 15 to 20 weeks’ foetuses were obtained from the MRC-Wellcome Trust Human Developmental Biology Resource (HDBR), UCL, with ethical approval (University College London, UCL, site REC reference: 18/LO/0822 – IRAS project ID: 244325 and Newcastle site REC reference: 18/NE/0290 – IRAS project ID: 250012).

### Monolayer cell impedance

xCELLigence (Agilent) is a system that continuously measures impedance across a cell layer^68^. HBMEC form tight junctions and are the first cell surface SpN must cross to invade the CNS. HBMEC (hCMEC/d3, Merck were cultured in Endogro^TM^-MV complete media (Millipore) with the addition of 1 ng/mL human Fibroblast Growth Factor (Merck. Cells were seeded on Collagen I (50 µg/mL, Merck) coated xCELLigence E-plate 16 at 8000 cells/well and placed in the xCELLigence RTCA DP system. When the cells had formed tight junctions (reached static impedance), *S. pneumoniae* isolates with previously enumerated CFU counts were thawed, washed and added to the wells in triplicate at MOI of 1. Plates were returned to the xCELLigence machine for ongoing quantification of impedance. Data were collected and analysed by real time cell analysis (RTCA) software supplied by the manufacturer.

## Multicellular blood brain barrier model

A multi-cellular transwell model of the BBB was developed from an original model developed using HBMEC, and Astrocytes.^69,70^

Primary astrocytes were were isolated from foetal brain samples as previously described^69–71^, human brain vascular pericytes (HBVP, ScienCell), and human microglia (HMC3, ATCC) were thawed from stock and cultured in 75 ml flasks coated with 2 μg/cm^2^ poly-l-lysine (astrocytes and pericytes) in cell-specific media (Astrocytes in DMEM (Sigma) + 10% FBS, HBVP pericyte growth media (ScienCell) and HMC3 in EMEM + 10% FBS, all supplemented with 100 U/ml penicillin, 100 µg/ml streptomycin (ThermoFisher) at 37 °C 5% CO_2_. HBMEC/d3s were incubated in Endogro-MV (Millipore) + FBGF as previously. All cells were used between passages 3-8.

All components for the BBB scaffold were sterilised in an autoclave at 100°C The apical surface of 6.5 mm diameter polycarbonate membrane transwells with 3 μm pores (Corning, New York) were coated with 50 μL of 150 μg/mL rat collagen-I solution and the basal surfaces coated with 50 μl at 2 μg/cm^2^ poly-l-lysine. Transwells were inverted and a sterilised section of rubber tubing was applied to the rim of the basal membrane. HBVP and astrocytes were passaged, counted and combined in a 150 μl aliquot containing 10,000 HBVP cells and 50,000 astrocytes per transwell. Cells were seeded on each basal transwell membrane surface and incubated for 4 hours at 37 °C 5% CO_2_ with regular media top up to prevent drying out. HBMEC/d3s were passaged, counted and diluted to 1.66×10^5^ cells/ml. The transwells were then righted, 150ul of hBEMC/d3s added to the apical surface of the transwell and gently lowered into 750uL of mixed media containing 50% each of Pericyte media and supplemented Endogro. The multi cellular transwell constructs were incubated at 37 °C 5% CO_2_ for up to 5 days. Permeability of the transwell model was assessed using Dextran diffusion as previously described.^70^ Briefly, Rhodamine B-labelled Dextran (Sigma) was diluted in 50% Endogro/Pericyte media to 0.5 mg/ml and a standard curve of 8x 1:2 dilutions was generated. 150ul media was removed from the apical chamber of each transwell and replaced with media containing Dextran-Rhodamine B. The transwells were incubated for 4 hours 37 °C 5% CO_2._ 100ul media was removed from the basal chamber of each transwell and fluorescence quantified using the Synergy Biotek2 plate reader (Agilent). Concentrations of Dextran-Rhodamine B were calculated against the standard curve. Cellular constructs where <20% Dextran was detected in the basal chamber compared to a blank transwell were deemed impermeable and used for subsequent experiments.

In parallel, HCM3 were seeded in 24-well plates and incubated for 24 hours in microglia specific media (Millipore) prior to BBB model infection. On the day of the experiment, this media was removed from the HMC3 cells, and replaced with 750 ul of ‘BBB media’ containing 1:1:1:1 mixture of Endogro/pericyte/astrocyte/microglia media.

The transwells containing the selected cellular constructs were placed into the 24-well plate containing microglial cells and incubated at 37°C at 5% CO_2_ while experimental conditions were prepared.

### Bacterial transmigration across multi-cellular BBB model

*S. pneumoniae* serotype 1 strains were incubated in THY + 0.5% yeast extract to mid-log phase, enumerated using CFU counting and frozen in 1 ml aliquots containing 80% glycerol. –Δ*Sp_1801-5*, Δ*bgaA* and Δ*ply*^48^ were incubated in media containing 0.1 mg/ml spectinomycin to inhibit growth of bacteria without the gene-deleted antibiotic cassette. Aliquots were thawed, washed and bacteria re-suspended in ‘BBB media’ at a MOI of 1. Earlier work indicated MOI higher than this resulted in rapid destruction of human cells. 150 ul of media was removed from the apical surface of the transwell and replaced with infected media. Each strain was tested in triplicate for each experiment. Bacterial counts at baseline in each media were enumerated using CFU plating as previously. The basal chamber was sampled for bacterial growth at hourly intervals for the first 4 hours, then again at 24 hours. 100 ul of media was removed from the basal chamber and plated in 1:10 dilutions to enumerate CFU Fresh media was placed in the basal chamber after each aspiration. Trans-endothelial electrical resistance (TEER) across each transwell insert was measured hourly for the first 8 hours and then at 24 hours.^72^

### Zebrafish embryo model

Pneumococcal injection stocks for zebrafish experiments were prepared by growing the cells in C+Y medium until an OD595 of 0.3 and then stored at –70 °C in medium with 20% glycerol. Before injection, bacteria were suspended in sterile PBS + 1% amaranth solution.

Adult wild-type zebrafish (Tupfel long fin line) were maintained at 26 °C in aerated tanks with a 10/14h dark/light cycle. Zebrafish embryos were collected within the first hours post fertilization (hpf) and kept at 28 °C in E3 medium (5.0 mM NaCl, 0.17 mM KCl, 0.33 mM CaCl·2H2O, 0.33 mM MgCl_2_·7H_2_O) supplemented with 0.3 mg/L methylene blue. At 1 day post fertilization (dpf) zebrafish were mechanically dechorionated.

Zebrafish were infected at 2 dpf by microinjection of 1500 for survival experiments or 2000 CFU for microscopy experiments of wild type or knockout mutant *S. pneumoniae* in the hindbrain vehicle.^73^ After infection zebrafish were kept in 6-well plates at 28 °C. For survival experiments, zebrafish were monitored at fixed time points until 72 hours post infection (hpi). Survival experiments were performed with 20 animals per group in quadruplicate.

To determine neutrophil infiltration of the cerebral ventricles live images were acquired with the Leica DMI6000 microscope. Tg(mpo:EGFP) zebrafish embryos were infected with wild-type or Δ*bgaA* strain through hindbrain injection and imaged at 3, 4 and 5 hours after infection. Embryos were embedded in 1.5 % low-melting-point agarose dissolved in egg water (60 μg/mL sea salts (Sigma-Aldrich; S9883) in MiliQ) in a µ-Slide 8 Well (Ibidi; 80826) immediately after injection and kept at 28°C during imaging. ImageJ software was used to process images and the number of neutrophils in the cerebral ventricle were counted in a blinded fashion by two independent researchers. Nine zebrafish were used per group and experiments were performed in triplicate.

To determine bacterial load after imaging, zebrafish were individually homogenized with zirconium balls in the MagNA lyser (Roche) instrument. The number of CFU per strain were determined by serial dilution of homogenates on COS plates containing COBA supplement (containing colistin sulphate 10 ug/ml and oxolinic acid 5 ug/ml). All procedures involving zebrafish embryos were performed according to local animal welfare regulations.

## Dutch cohort study

We studied the role of *bgaA* or *Sp_1801-5* in a Dutch prospective nationwide study of bacterial meningitis, the Meningene study. This study included patients ≥16 years who are listed via bacterial monitoring by the Netherlands Reference Laboratory for Bacterial Meningitis (NRLBM). This lab receives samples from both CSF and blood from around 85% of the Dutch bacterial meningitis patients. Detailed methodology for patient selection and inclusion has previously been published^33^. In summary, the NRLBM provided daily updates of hospitals in which patients with bacterial meningitis had been admitted in the preceding days and names of treating physicians. Physicians were informed by telephone about the study or could contact Meningene investigators themselves to include patients. Patients or their legal representatives were given written study information and asked for written informed consent. Baseline, admission, treatment and outcome data was collected by the treating physician using an online case record form.

Pneumococcal meningitis was defined as a CSF culture positive for *S. pneumoniae* or a combination of a blood culture, CSF PCR or CSF antigen test positive for *S. pneumoniae* with CSF chemistry indicative of bacterial meningitis according to the criteria defined by Spanos *et al*. (glucose <1.9 mmol/L, CSF-blood glucose ratio <0.23, CSF protein >2.2 g/L, CSF leukocyte count >2000 per mm3, or more than 1180 polymorphonuclear leukocytes per mm3).^74^ Patients with hospital acquired bacterial meningitis, recent neurosurgery (≤1 month), recent neurotrauma (≤1 month), or with a neurosurgical device in place were excluded.

Pathogens were stored at –80 °C in the NRLBM upon receipt. For DNA extraction, isolates were re-cultured from frozen stocks on blood agar plates. Sequencing was performed using multiplexed libraries on the Illumina HiSeq platform to produce paired end reads of 100 nucleotides in length (Illumina, San Diego, CA, USA).

The infecting strain was routinely genotyped in 1025 pneumococcal meningitis patients included in the Meningene. To determine if the *bgaA* or *Sp_1801-05* loci were present we performed a BLAST analysis using all known loci from PubMLST. *BgaA* or *Sp_1801-05* was classified as present if a locus was found with at least 99% similarity to a PubMLST locus. Clinical characteristics were compared between patients infected with strains with or without a *bgaA* or *Sp_1801-05* locus.

### Murine transnasal brain infection model

*S. pneumoniae* deletion mutant strains were tested in a mouse model of nasopharynx-to-brain translocation model previously described.^55^ Groups of 5 mice anaesthetized with 2.5% isoflurane and intranasally inoculated with a 10 μl suspension containing 10^8^ CFU of wild-type, Δ*bgaA* or Δ*Sp_1801-05* strains. At predetermined time points, mice were culled and the CFU were determined at 0, 3, 7, and 10 days post-inoculation in tissue samples including the nasopharynx (NP), olfactory bulb (OB), olfactory epithelium (OE) and Brain (Br) as previously described.^55^ Blood was also checked for CFU at 3, 7 and 10 days post-infection, and no bacteria were detected in blood at any timepoint.

## Declaration of Interests

All authors have no conflicts of interest to declare

## Funding

This study was funded by a Clinical Lecturer Starter Grant from the Academy of Medical Sciences (UK) and Wellcome Trust Institutional Strategic Support Funding (ISSF) with the Robin Weiss Fund to EW. The Bundles for Adult Meningitis (BAM) study was funded by a PhD Fellowship in Global Health to EW from the Wellcome Trust (089671/B/09/Z). Additional funding included a Postdoctoral Clinical Research Fellowship to EW from the Francis Crick Institute. The Malawi-Liverpool-Wellcome Trust Clinical Research Programme is supported by a core grant from the Wellcome Trust (101113/Z/13/Z). The laboratory work was undertaken in part at UCLH/UCL who received a proportion of funding from the National Institute for Health Research University College London Hospitals Department of Health’s NIHR Biomedical Research Centre. ECW, RH and JSB are supported by the Centre’s funding scheme. RJW, AP and EW were supported by the Francis Crick Institute which receives funding from Wellcome (CC2112), Cancer Research UK (FC2112) and UK Research and Innovation: Medical Research Council (FC2112). The human foetal material was provided by the Joint MRC/Wellcome Human Developmental Biology Resource (www.hdbr.org) (project#200511). RSH is a NIHR Senior Investigator.

Additional laboratory work at UCL and LSHTM was funded by an Investigator award to JSB and BW from the Wellcome Trust.

For the purposes of open access, the authors have applied a CC-BY public copyright to any author-accepted manuscript arising from this submission.

The funders of the study had no role in study design, data collection, data analysis, data interpretation, or writing of the report. The corresponding author had full access to all the data and the final responsibility to submit for publication.

## Author contributions

Conception or design of the work: ECW, JAGA, AK, MY, RK, DvdB, JWV, BW, RJW, RSH, JSB

Data collection: ECW, JAGA, VST, ERS, GE, AP, MY, AT, RK, RA

Data analysis and interpretation: ECW, JAGA, BW, AK, MY, JSB

Writing and editing: EW, JSB, JAGA, MY, VST, RK, DvdB, RJW, JWV, AK, RSH,JSB

Final approval of the version to be published: all authors

## Acknowledgements

The authors would like to thank the study patients and guardians, the Bundles for Adult Meningitis (BAM) research team, clinical and laboratory staff at the Queen Elizabeth Central Hospital and Malawi-Liverpool-Wellcome Trust Clinical Research Programme in Blantyre Malawi for support given during the study and Professor Mike Levin and Dr Victoria Wright of Imperial College UK for support at the early stages of the project and donation of the PAXgene tubes to collect CSF. We would like to thank Professor Judith Breuer and the staff of the Pathogen Genomics Unit at University College London for their assistance with library preparation and RNA sequencing. The authors acknowledge the use of the UCL Legion High Performance Computing Facility (Legion@UCL), and associated support services, in the completion of this work.

## Tables

**Table 1:** Summary of patient and sample characteristics.

**Table 2:** Highly expressed gene loci during human meningitis

## Supplementary tables / figures (10 total maximum)

**Supplementary Table 1:** Comparison of the pneumococcal transcriptome in human CSF to infection-mimicking conditions in the Pneumoexpress D39 culture transcriptomic model^75^

**Supplementary table 2:** Representation of regulons (described in RegPrecise^42^ for strain TIGR4) amongst highly transcribed genes from the human meningitis RNAseq data

**Supplementary Table 3:** In silico functional predictions for the corresponding protein for genes within the Sp_1801-1805 operon

**Supplementary Table 4**: Genome associations and outcomes in human meningitis using genome data from 1144 strains isolates from European cases of meningitis and the associated clinical data^9^.

## Supplementary Figure Legends

**Supplementary Figure 1: Kegg analysis of over-expressed metabolic pathways in *S. pneumoniae* derived from the human meningitis pneumococcal transcriptome**. Highly enriched KEGG metabolic pathways in pneumococcal meningitis, overlayed onto the entire pneumococcal metabolic network available in KEGG. Nodes represent individual genes, lines represent metabolic pathways connecting genes within a metabolic network/pathway. Green networks are upregulated, red networks are down-regulated. For a list of individual metabolic pathways see Table 2A.

**Supplementary Figure 2** – Visualisation of protein structure predictions for Sp_1804 generated by HMMER and rendered in Phyre2

**Supplementary Figure 3:** Composition of a four cell *in vitro* transwell model of the Blood Brain Barrier.

**Supplementary data file:** Alignment, mapping, sequening and transcriptome data. Differential gene expression of *S. pneumoniae* in vitro between CSF, CSF + Neutrophils and THY conditions.

## Notes

### Competing Interest Statement

The authors have declared no competing interest.

