## Supplementary Tables for "The *Streptococcus pneumoniae* transcriptome in patient cerebrospinal fluid identifies novel virulence factors required for meningitis"

**Supplementary Table 1: Comparison of the pneumococcal transcriptome in human CSF to infection-mimicking conditions in the Pneumoexpress D39 culture transcriptomic model[69].**

Patient pneumococcal transcriptomes in CSF were re-mapped against D39 and analysed in the D39 transcriptomic model, testing for clusters of co-expressed genes across the patient samples and model systems. Cluster expression indicates overlap between highly expressed genes in the human pneumoccocal transcriptome and each individual condition. Enrichment score indicates the proportion of shared genes across the cluster.

| Pneumoexpress conditions | Cluster expression (CPM) | Enrichment score |
|---|---|---|
| **LMC** | 2227 | 4.85 |
| **BMC** | 2826 | 6.21 |
| **CSF mimicking condition** | 2479 | 5.42 |
| **Fever** | 3103 | 6.84 |
| **C+Y complete media** | 3884 | 8.64 |
| **competence, 3 min** | 3368 | 7.45 |
| **competence, 10 min** | 2297 | 5.01 |
| **competence, 20 min** | 3724 | 8.27 |
| **Infection A549, 0 mpi** | 3385 | 7.49 |
| **Infection A549, 30 mpi** | 4258 | 9.52 |
| **Infection A549, 60 mpi** | 4273 | 9.56 |
| **Infection A549, 120 mpi** | 3589 | 7.96 |
| **infection, A549, 240 mpi** | 3570 | 7.91 |

LMC = Lung mimicking condition, BMC = Blood mimicking condition.

**Supplementary Table 2:** Representation of regulons (described in RegPrecise[40] for strain TIGR4) amongst highly transcribed genes from the human meningitis RNAseq data.

| Regulator | Role | Number of genes highly expressed in human meningitis / total regulon |
|---|---|---|
| MntR: Sp_1638 | Manganese homeostasis | 4 / 4 |
| BirA: Sp_1900 | Biotin synthesis | 1 / 1 |
| FabT: Sp_0416 | Fatty acid biosynthesis | 12 / 12 |
| RNA - L21_leader | Ribosome biogenesis | 3 / 3 |
| NrdR: Sp_1713 | Deoxyribonucleotide biosynthesis | 3 / 4* |
| AgaR: Sp_0058 | N-acetylgalactosamine utilisation | 5 / 12 |
| CodY: Sp_1584 | Amino acid metabolism | 8 / 42 |
| Rex: Sp_1090 | Energy metabolism | 2 / 11 |
| CcpA: Sp_1999 | Global catabolite repression | 14 / 171 |
| RNA – PyrR | Pyrimidine metabolism | 1 / 9 |

* 4 extra genes from the regulon missing from meningitis RNAseq transcripts (Sp_0203-5 and 1178)
** 1 extra gene from the regulon missing from meningitis RNAseq transcripts (Sp_0415)

**Supplementary Table 3:** In silico functional predictions for the corresponding protein for genes within the Sp_1801-1805 operon

| Gene | Protein analysis software | | | | | | |
|---|---|---|---|---|---|---|---|
| | **BLASTp identification** | **InterPro** | **PANNZER** | **DMPFold** | **Phyre2** | **HMMER vs UniProt / PFAM** | **HHblits** |
| SP_1801 | GlsB/YeaQ/YmgE family stress response membrane protein https://www.ncbi.nlm.nih.gov/protein/WP_000907510.1?report=genbank&log$=prottop&blast_rank=2&RID=6PVG7V0M013 | Transglycosylase-associated protein | GlsB | none | cryo-em structure of euglena gracilis mitochondrial atp synthase,2 membrane region | Uncharacterized protein https://www.uniprot.org/uniprot/A0A0H2URM3 Transglycosylase associated protein | YeaQ/YmgE, transglycosylase membrane protein |
| SP_1802 | Hypothetical / alkaline shock response membrane anchor protein AmaP https://www.ncbi.nlm.nih.gov/protein/WP_000030213.1?report=genbank&log$=prottop&blast_rank=3&RID=6PVUMZH5016 | none | Alkaline shock response membrane anchor protein AmaP | http://bioinf.cs.ucl.ac.uk/psipred/&uuid=ddea2528-d7e9-11ea-8d55-00163e100d53 | | Alkaline shock membrane protein https://www.uniprot.org/uniprot/B2ISG1 | Alkaline shock response membrane anchor protein AmaP |
| SP_1803 | DUF2273 domain-containing protein, or putative lipoprotein https://www.ncbi.nlm.nih.gov/protein/WP_050239154.1?report=genbank&log$=prottop&blast_rank=2&RID=6PW4JBME016 | Unknown function protein DUF2273 | DUF2273 domain-containing protein (Fragment) | none | none | Putative lipoprotein / Small integral membrane protein (DUF2273) | https://www.uniprot.org/uniref/UniRef100_A0A064C068 |
| SP_1804 | Asp23/Gls24 family envelope stress response protein https://www.ncbi.nlm.nih.gov/protein/WP_000064115.1?report=genbank&log$=prottop&blast_rank=1&RID=6PWBEFFK013 | Alkaline shock protein Asp23 | General stress protein | http://bioinf.cs.ucl.ac.uk/psipred/&uuid=ca0c47ac-d7e9-11ea-b425-00163e100d53 | Suppl. Fig. 2 | Gls24 protein https://www.uniprot.org/uniprot/B1I7R1 | General stress protein, Gls24 family |
| SP_1805 | CSBD family protein https://www.ncbi.nlm.nih.gov/protein/WP_000109957.1?report=genbank&log$=prottop&blast_rank=1&RID=6PWYZY4Y01N | CsbD stress response protein | CsbD domain-containing protein | none | Fold:SAM domain-like Superfamily: YjbJ | CsbD http://www.uniprot.org/uniprot/A0A2N6SHC1_9BACL | CsbD family protein |

**Supplementary Table 4**: Genome associations and outcomes in human meningitis using genome data from 1144 strains isolates from European cases of meningitis and the associated clinical data[9].

| Variable | *bgaA* positive | *bgaA* negative | P value (Mann-Whitney U test) |
|---|---|---|---|
| **Demographics** | | | |
| **Age (years)** | 61 (49-70) | 57 (50-67) | 0.09 |
| **Female sex** | 465/901 (52%) | 66/124 (53%) | 0.40 |
| **Immunocompromised state** | 237/900 (26%) | 29/124 (23%) | 0.28 |
| **Diabetes Mellitus** | 117/888 (13%) | 11/123 (9%) | 0.12 |
| **Alcoholism** | 54/896 (6%) | 8/123 (7%) | 0.48 |
| **Otitis** | 316/857 (37%) | 35/119 (29%) | 0.07 |
| **Sinusitis** | 124/838 (15%) | 16/117 (14%) | 0.44 |
| **Pneumonia** | 104/871 (12%) | 17/121 (14%) | 0.30 |
| **Symptoms on Admission** | | | |
| **Coma on admission** | 246/898 (27%) | 31/124 (25%) | 0.33 |
| **Glasgow Coma Scale on Admission** | 10 (8-13) | 10 (8-12) | 0.89 |
| **Heartrate on admission** | 100 (86-115) | 100 (88-112) | 0.91 |
| **Laboratory results on admission** | | | |
| **Leukocytes CSF** | 2625 (542-7955) | 4330 (975-9725) | 0.03 |
| **Leukocytes blood** | 17 (12-23) | 18 (13-24) | 0.20 |
| **CSF:blood glucose ratio** | 0.03 (0.01-0.20) | 0.03(0.01-0.19) | 0.81 |
| **Clinical course** | | | |
| **Mortality** | 202/901 (22%) | 16/124 (13%) | 0.008 |
