## Supplementary Figure 1 for "The *Streptococcus pneumoniae* transcriptome in patient cerebrospinal fluid identifies novel virulence factors required for meningitis"

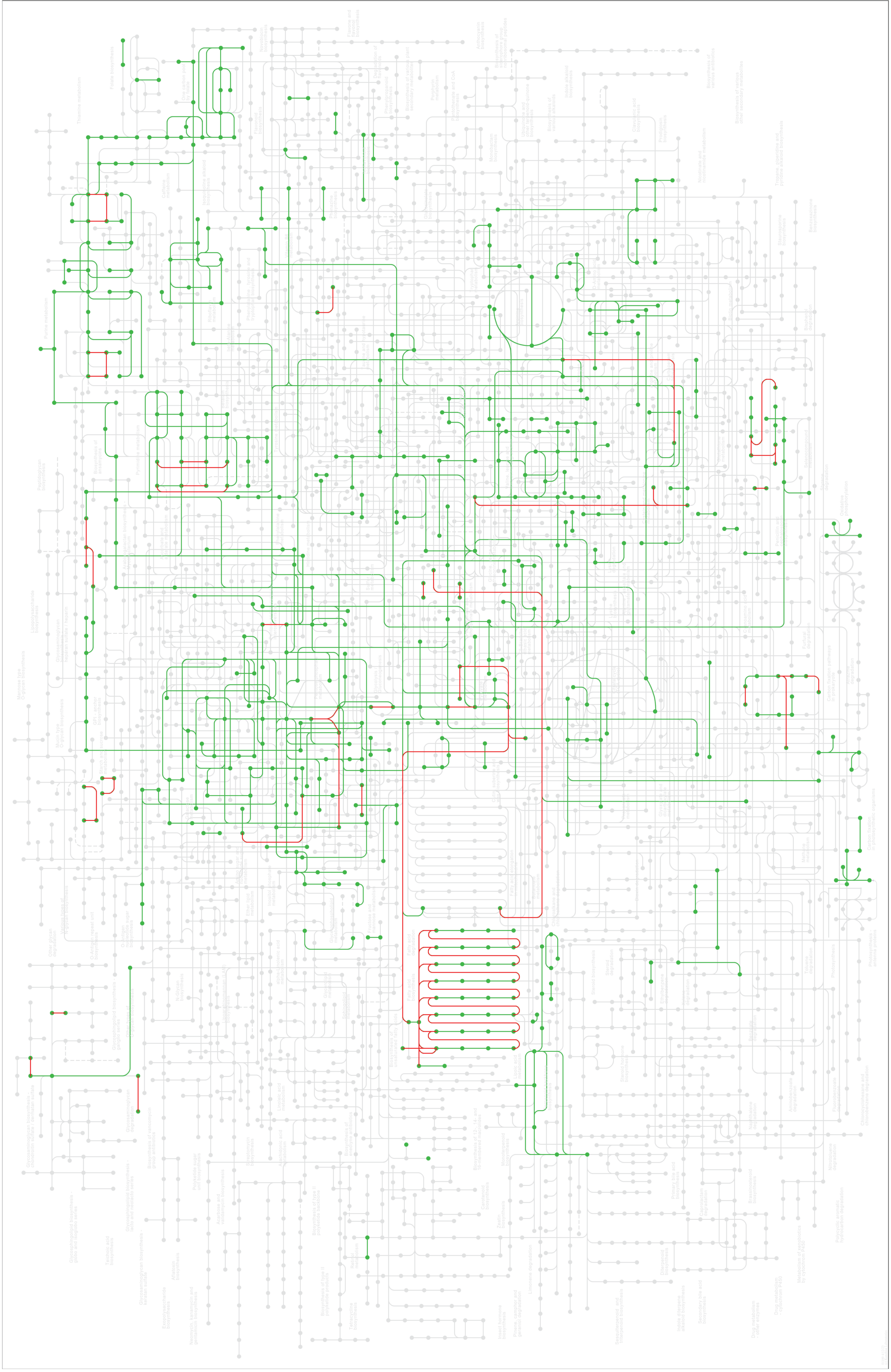

**Supplementary Figure 1: KEGG pathway analysis of over-expressed metabolic pathways during pneumococcal meningitis.**

Highly enriched KEGG metabolic pathways in pneumococcal meningitis, overlayed onto the entire pneumococcal metabolic network in KEGG. Nodes represent individual genes, lines represent metabolic pathways connecting genes within a metabolic network/pathway**.** Green networks are upregulated, red networks are down regulated. For a list of individual metabolic pathways see table 2A
