## Supplementary FIgure 2 for "The *Streptococcus pneumoniae* transcriptome in patient cerebrospinal fluid identifies novel virulence factors required for meningitis"

**A**

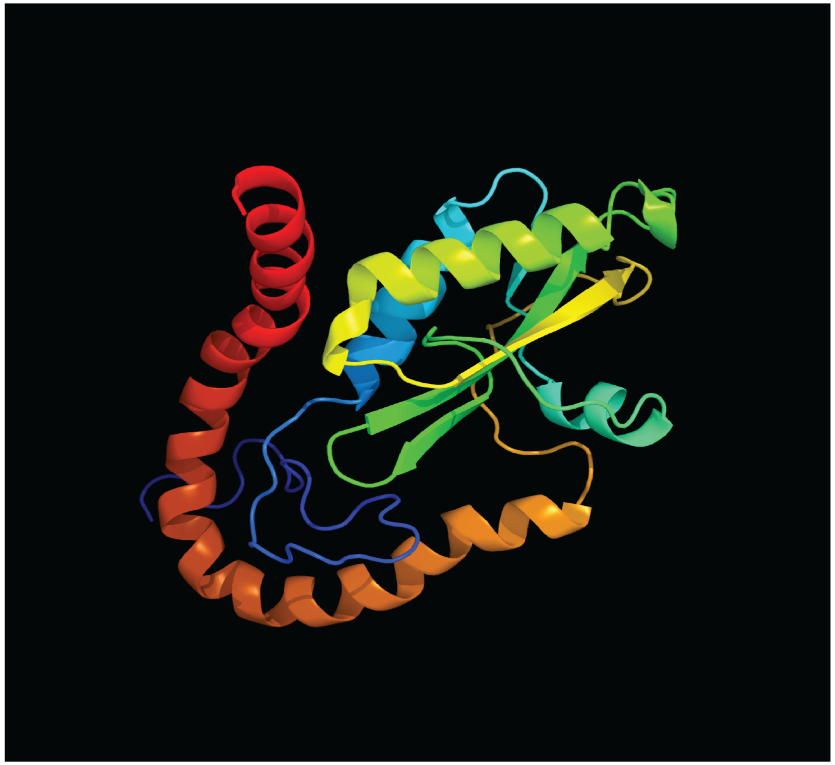

**Supplementary figure 2: Predicted protein structure for (A) Sp_1804**

Visualisation of protein structures generated using Phyre2 after sequence analysis in HMMER. Colour indicates
