## Supplementary Figure 3 for "The *Streptococcus pneumoniae* transcriptome in patient cerebrospinal fluid identifies novel virulence factors required for meningitis"

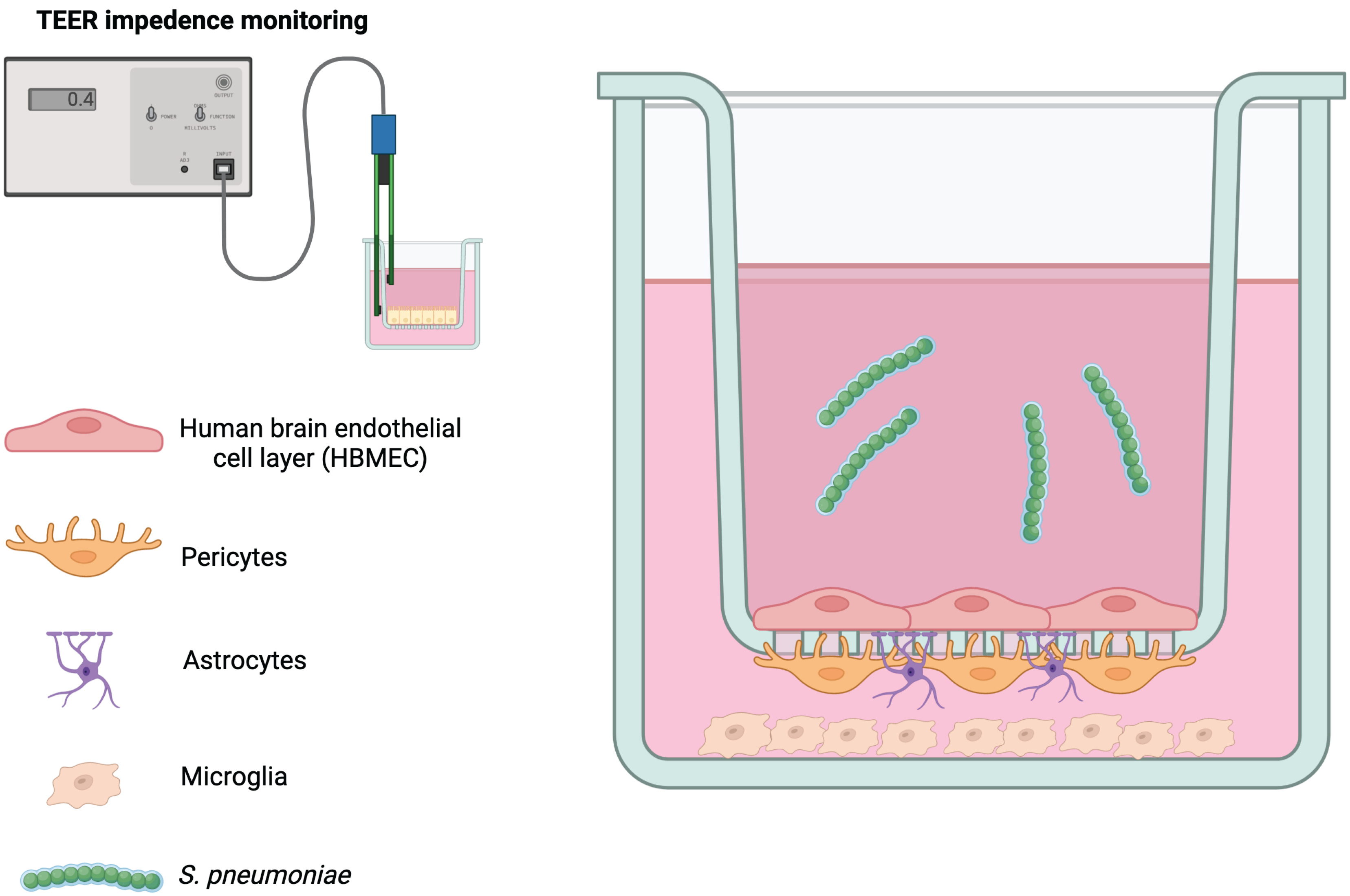


**Supplementary figure 3: Composition of a 4-cell *in vitro* transwell model of the blood brain barrier (BBB)**
